# Enhanced transcriptional strength of HIV-1 subtype C minimizes gene expression noise and confers stability to the viral latent state

**DOI:** 10.1101/2022.08.04.502743

**Authors:** Sreshtha Pal, Vijeta Jaiswal, Narendra Nala, Udaykumar Ranga

**Author notes:** Current Address: Synthetic Biology Group, Institute of Bioinformatics and Applied Biotechnology, Bangalore, Karnataka, India (Vijeta Jaiswal). Corresponding Author: Udaykumar Ranga (UR).

## Abstract

The stochastic fluctuations in gene expression emanating from HIV-1 long terminal repeat (LTR), amplified by the Tat positive feedback circuit, determine the choice between viral infection fates: active transcription (ON) or transcriptional silence (OFF). The emergence of several transcription factor binding site (TFBS) variant strains in HIV-1 subtype C (HIV-1C), especially those containing the duplication of NF-κB motif, mandates the evaluation of the effect of enhanced transcriptional strength on gene expression noise and its influence on viral fate-selection switch. Using a panel of subgenomic LTR-variant strains containing varying copy numbers of the NF-κB motif (ranging from 0 to 4), we employed flow cytometry, mRNA quantification, and pharmacological perturbations to demonstrate an inverse correlation between promoter strength and gene expression noise in Jurkat T-cells and primary CD4^+^ T-cells. The inverse correlation is consistent in clonal cell populations, at constant intracellular concentrations of Tat, and when NF-κB levels were regulated pharmacologically. Further, we show that strong LTRs containing at least two copies of the NF-κB motif in the enhancer establish a stabler latent state and demonstrate rapid latency reversal than weak LTRs containing fewer motifs. An engineered LTR containing three copies of the C-κB motif (CCC), an element unique for HIV-1C, demonstrated significantly higher levels of gene expression noise compared to the canonical HHC-LTR or two other engineered LTRs containing three copies of the H-κB (HHH) or F-κB (FFF) motif. This result suggests the indispensable nature of the C-κB motif for HIV-1C despite higher-level gene expression noise. We also demonstrate a cooperative binding of NF-κB to the motif cluster in HIV-1C LTRs containing two, three, or four NF-κB motifs (H = 2.61, 3.56, and 3.75, respectively). The present work alludes to a possible evolution of HIV-1C LTR towards gaining transcriptional strength associated with attenuated gene expression noise with implications for viral latency.

**Author Summary:** Over the past two consecutive decades, HIV-1C has been undergoing directional evolution towards augmenting the transcriptional strength of the LTR by adding more copies of the existing TFBS by sequence duplication. Additionally, the duplicated elements are genetically diverse, suggesting broader-range signal receptivity by variant LTRs. HIV-1 promoter is inherently noisy, and the stochastic fluctuations in gene expression of variant LTRs may influence the ON/OFF latency decisions. The evolving NF-κB motif variations of HIV-1C offer a powerful opportunity to examine how the transcriptional strength of the LTR might influence gene expression noise. Our work here shows that the augmented transcriptional strength of HIV-1C LTR leads to a concomitantly reduced gene expression noise, consequently leading to stabler latency maintenance and rapid latency reversal. The present work offers a novel lead towards appreciating the molecular mechanisms governing HIV-1 latency.

## Introduction

Gene expression noise is pervasive and universal; therefore, biological processes, especially those regulating essential functions, attempt to keep the noise to a minimum (1,2) (PMID: 16369570, 23953111). Various mechanisms minimizing gene expression noise have been identified (3–5) (PMID: 15124029, 20164922, 34625470). The emerging single-cell analytical techniques permit the appreciation of the significance of gene expression noise to biological processes (6) (PMID: 19416069). For instance, studies in yeast demonstrated that the augmented phenotypic variations in isogenic populations could be ascribed to gene expression noise (7,8) (PMID: 18362885, 17189188). Thus, gene expression noise could enable populations to adopt a bet-hedging strategy to optimize survival fitness in a fluctuating environment. From an evolutionary point of view, gene expression noise can diversify a range of phenotypes from a single genotype (9) (PMID: 19401676). Driving the fate decisions in a fluctuating environment, also called ‘stochastic cell-fate specification’, represents the most important functional role ascribed to gene expression noise that gained maximum focus (10) (PMID: 21414483). A prominent example is *Bacillus subtilis,* which chooses between dormant and competent states by stochastically altering the levels of a key regulatory protein (11) (PMID: 17569828).

Likewise, viruses that encounter fluctuating environments in a cell must make quick fate choices to thrive in the host. In such a case, fluctuations in gene expression can play a pivotal role in driving fate decisions and gaining a fitness advantage. Decision-making between two viral fate states: lytic and lysogeny phases, in the *λ*-phage, is the outcome of noise in the expression of two regulatory proteins Cro and CI (12,13) (PMID: 9691025, 10659856). Stochastic fluctuations at the promoters P_R_ and P_RM_ are reflected at the levels of the gene products Cro and CI proteins, respectively. A critical increase or decrease in Cro and CI levels could drive the decision-making between the lytic and lysogeny fates.

Among animal viruses, HIV-1 is believed to use stochastic fluctuations of the expression of Tat, the master regulatory protein of the virus, to drive the decision-making between active replication and viral latency (14) (PMID: 16051143). Following infection of target CD4^+^ T-cells, HIV-1 integrates into the host genome at random sites and chooses between active replication and viral latency (15) (PMID: 22762018). The ‘latent’ state of the virus offers the ultimate challenge to a functional cure as the available cure strategies cannot eradicate the latent viral reservoir.

A significant level of controversy surrounds the mechanisms underlying ‘HIV-1 latency’. One school of thought posits that HIV-1 latency is an ‘epiphenomenon’, where latency is deterministically regulated by the transition of an infected T-cell from an activated state to a resting phenotype (16,17) (PMID: 22229121, 23284080). Alternative studies suggest that the decision-making is hardwired into the virus, i.e., the intrinsic mechanisms randomly regulating viral transcription and latency reversal (18,19) (PMID: 24243014, 25723172). Notably, the viral master transcription regulatory circuit (MTRC) consisting of the cis-acting LTR and the trans-acting Tat positive feedback loop, along with stochastic fluctuations of viral transcription, orchestrate the switch between the ON/OFF fates (14,20) (PMID: 16051143, 18344999). Thus, gene expression noise could be a critical factor influencing the ON/OFF decision-making of HIV-1, especially in the absence of activation. Of note, gene expression from the HIV-1 promoter is inherently noisy relative to mammalian promoters (21,22) (PMID: 20409455, 23064634), which could be ascribed to the complex promoter architecture. HIV-1 promoter houses binding sites for several families of transcription factors within a span of ~450 base pairs (23) (PMID: 10637316). Given the densely packed organization, several of the TFBS are tandemly arranged and tend to overlap with each other. Additionally, the NF-κB and Sp1 binding sites are present in multiple copies and arranged tandemly.

There exists no consensus regarding a correlation between the transcriptional strength of a eukaryotic promoter and gene expression noise; while several studies demonstrate an indirect correlation between the two (24–26) (PMID: 15166317, 23565060, 20159560), others reported a direct correlation (27–31) (PMID: 16715097, 16699522, 17048983, 20185727, 25030889). Broadly, most of these studies underline the significance of the promoter architecture being a crucial parameter influencing the gene expression noise of a promoter. Thus, the high-density arrangement of TFBS belonging to several transcription factor families and the presence of multiple copies of specific TFBS responsive to excitatory cellular activation signals could underlie the significantly higher-level transcriptional noise of the HIV-1 LTR, in turn influencing the ON/OFF decision-making.

If gene expression noise of the LTR is a function of the TFBS profile, then functional inactivation of specific sites may alter the ON/OFF decision-making. A few studies inactivated selected TFBS of the LTR and demonstrated that the inactivation of the TATA box and the Sp1 elements, but not the NF-κB motifs, modulated gene expression noise (32,33) (PMID: 19132086, 23874178). Likewise, manipulating the Tat-feedback strength is also expected to change viral latency decisions. For example, tweaking the Tat positive feedback strength modulates the duration of fluctuations in Tat levels, in turn, modulating viral latency decisions (20) (PMID: 18344999). Thus, individual TFBS in the LTR can exert a modulatory effect on viral transcription.

Over the years, our laboratory monitored the evolution of HIV-1C in India and identified two major hotspots of genetic variation in the viral genome - in the LTR and p6-Gag (34–38) (PMID: 15184461, 23132857, 24960272, 29773649, 34899661). The variations in the LTR are centered around the creation of additional copies of certain TFBS by sequence motif duplication. A recent study from our laboratory identified the emergence of at least nine different promoter variant viral strains of HIV-1C containing duplications of TFBS belonging to TCFα/LEF, RBE III, and/or NF-κB (36) (PMID: 34899661). Of note, the LTR variations are characterized by not only copy number difference but also sequence variation and the co-duplication of the TFBS motifs. For example, unlike the canonical HIV-1 subtype B (HIV-1B) LTR containing two genetically identical copies of the NF-κB motif (HH), the canonical HIV-1C LTR contains three copies of the NF-κB motif (HHC). A subset of HIV-1C LTR variant strain also contains four copies (FHHC). Importantly, the additional NF-κB motifs are genetically distinct from the canonical H-κB motif and even from each other. Thus, the 4-κB LTR of HIV-1C contains three genetically different NF-κB binding sites in the enhancer. Notably, a direct correlation exists between the NF-κB binding sites in the LTR and the transcriptional strength. Further, the 4-κB viral strains dominated the 3-κB strains under all experimental conditions and in natural infection (34) (PMID: 23132857). Significantly, we also demonstrated that enhanced transcriptional strength of the LTR leads to faster latency establishment (39) (PMID: 32669338).

Thus, the enhanced complexity of the LTR architecture of HIV-1C is expected to make viral purging more challenging on the one hand but likely to offer crucial leads into the molecular mechanisms governing viral latency on the other hand. To this end, using panels of subgenomic NF-κB motif copy number variant viral strains, we evaluated gene expression noise modulation and its impact on HIV-1C latency. We demonstrate an inverse correlation between NF-κB motif copy number and gene expression noise, potentially impacting viral latency maintenance.

## Materials and Methods

### Cell culture

Human embryonic kidney cells (HEK293T) were cultured in Dulbecco’s Modified Eagle Medium (DMEM; Catalog no. D1152; Sigma-Aldrich) supplemented with 10% fetal bovine serum (FBS; Catalog no. RM10435; HiMedia Laboratories, India), 2 mM glutamine (Catalog no. G8540; Sigma, St. Louis, MO), 100 U/ml penicillin G (Catalog no. P3032; Sigma, St. Louis, MO), and 100 g/ml streptomycin (Catalog no. S9137; Sigma, St. Louis, MO). Jurkat T-cells were cultured in Roswell Park Memorial Institute Medium (RPMI 1640; Catalog no. R4130; Sigma, St. Louis, MO) supplemented with 10% FBS, 2 mM glutamine, 100 U/ml penicillin G, 100 g/ml streptomycin. The peripheral blood mononuclear cells (PBMC) were isolated from 15 ml of fresh blood drawn from healthy donors by density gradient centrifugation using Histopaque-1077 (Catalog no. 10771; Sigma-Aldrich). Subsequently, primary CD4^+^ T-cells were negatively enriched using EasySep Human CD4^+^ T-cell isolation kit (Catalog no. 17952; Stem Cell Technologies) and cultured in RPMI supplemented with 10% FBS and 20 U/ml of Interleukin-2 (IL-2; Catalog no. H7041; Sigma, St. Louis, MO). All the cells were incubated at 37°C in the presence of 5% CO_2_.

### Construction of HIV-1 reporter panels

#### (i) Construction of the cLdGIT panel

The initial pLGIT (HIV-1B LTR-EGFP-IRES-Tat) vector (14) (PMID: 16051143) was a kind gift from Dr. David Schaffer (University of California, USA), which was modified in various successive steps to substitute Tat and LTR of HIV-1B origin to HIV-1C origin (BL4-3; GenBank accession no. FJ765005.1) (40) (PMID: 27194770); and EGFP to d2EGFP, a destabilized form of EGFP with a half-life of ~2 h (39) (PMID: 32669338). The resulting bicistronic vector cLdGIT HHC (HIV-1C LTR-d2EGFP-IRES-Tat) that encoded d2EGFP and HIV-1C Tat driven by HIV-1C LTR was used to study the stochastic nature of the ‘Mid’ population.

Chakraborty et al. modified the cLdGIT vector with three canonical NF-κB motifs (HHC) to a variant with four NF-κB motifs (FHHC), widely prevalent in the population (39) (PMID: 32669338). Using cLdGIT FHHC as the parental vector, inactivating point mutations were introduced sequentially to the FHHC NF-κB motif to generate a panel of vectors with a varying number of NF-κB binding motifs, ranging from 0 to 4 copies. First, using overlap PCR and cLdGIT FHHC as the template, LTR-fragments with inactivating point mutations at each subsequent NF-κB motif were amplified to generate OHHC (3-κB), OOHC (2-κB), OOOC (1-κB), OOOO (0-κB) LTR-fragments. We mutated the critical residues of the NF-κB motifs essentially required for NF-κB transcription factor binding (41) (PMID: 17869269), without ablating the NFAT binding sites. Unique RE sites were engineered into the panel vectors, downstream of the 3’ LTR using the outer reverse primer for the unique identification of each vector. The primer details and unique restriction enzyme sites corresponding to each variant viral strain are presented in Table 1. The amplified LTR-fragments were directionally cloned between XhoI and PmeI sites, present in the outer forward and outer reverse primers, respectively, thereby replacing the FHHC-LTR at the 3’ end. Thus, all the LTR-variant viral strains were genetically identical except in the number of NF-κB motifs in the LTR. The recombinant clones were identified using restriction digestion and validated using Sanger sequencing. EGFP expression of all cLdGIT variant viral strains was validated in HEK293T cells.

**Table 1.**
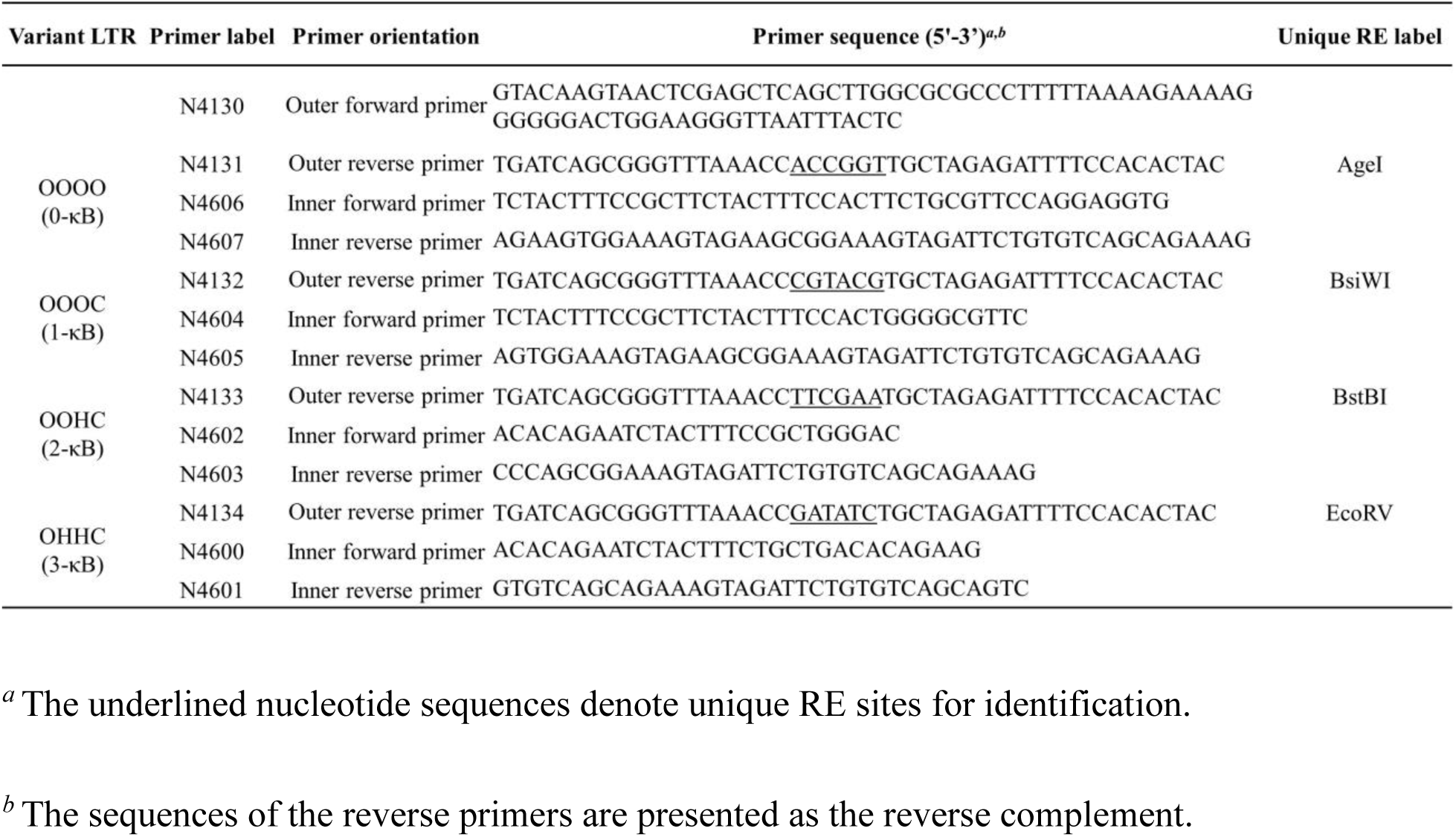
Primer sets used for site-directed mutagenesis of NF-κB motif copy-number variation in pcLdGIT FHHC backbone.

#### (ii) Construction of the cLdG NF-κB-variant panel

In the cLdG (HIV-1C LTR-d2EGFP) panel, unlike in the cLdGIT panel, the expression of only d2EGFP, but not that of Tat, was placed under the LTR. Using ApaI and XhoI, the ‘IRES-Tat’ region of the cLdGIT variant viral strains was excised, and the remaining backbone was self-ligated. EGFP expression of all the cLdG variant viral strains was validated in HEK293T cells.

#### (iii) Construction of a panel of viral strains containing homotypic clusters of NF-κB motifs in the LTR

Using the canonical cLdGIT HHC vector as the parental backbone, we generated a panel of LTR-variant viral strains containing three copies of one of the three genetically distinct NF-κB motifs (H, C, and F-κB sites), naturally present in HIV-1C. The LTR fragments of HHH, FFF, and CCC-κB motifs were amplified using an overlap-PCR. The amplified fragments were used to replace the HHC region of the parental cLdGIT HHC vector between XhoI and PmeI sites. Unique RE labels were introduced into each vector at the end of the 3’ LTR upstream of the PmeI site of outer reverse primers. The primer details and unique RE identification labels are summarised (Table 2). The recombinant clones were identified by restriction digestion and validated using Sanger sequencing. EGFP expression was confirmed in HEK293T cells.

**Table 2.**
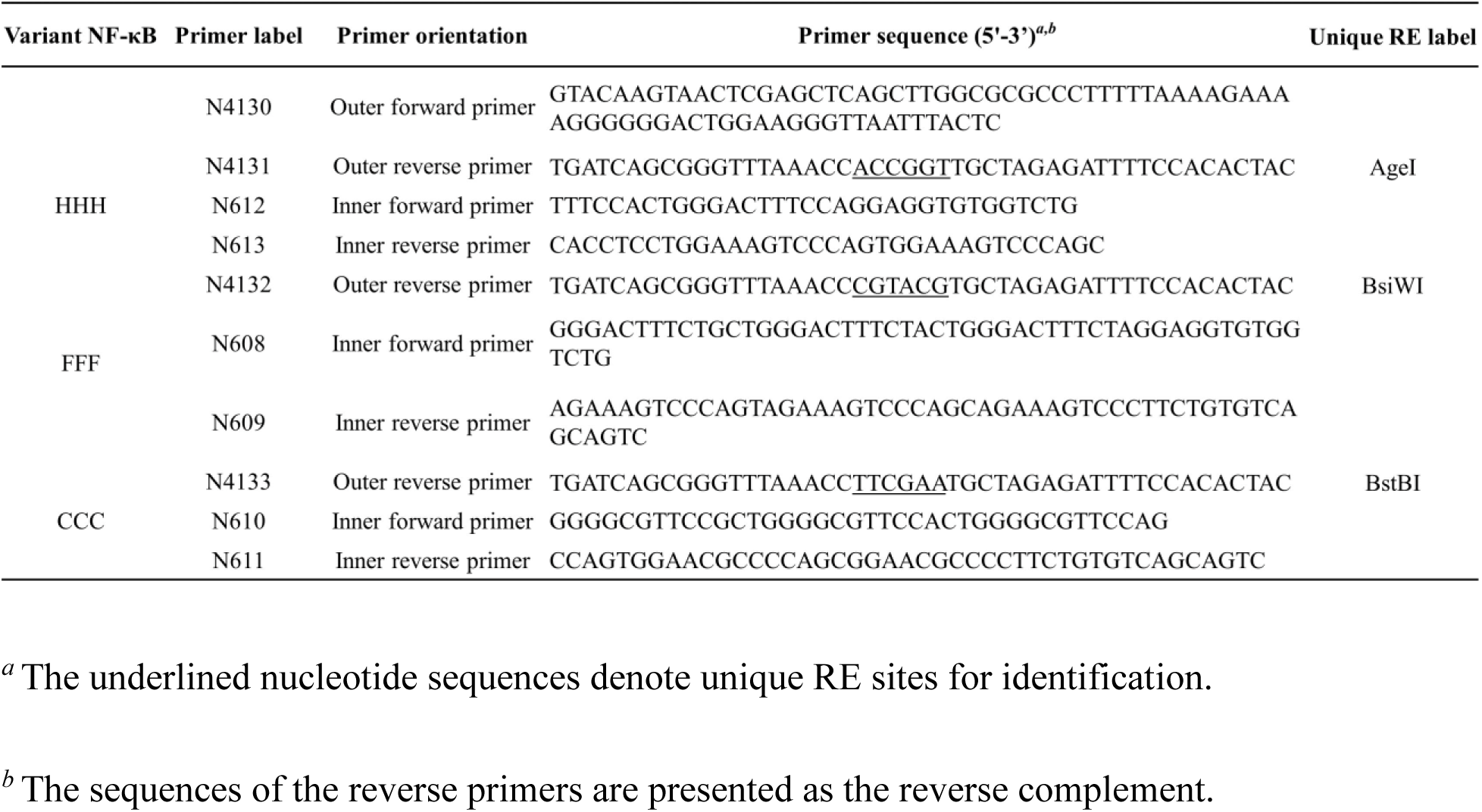
Primer sets used to generate cLdGIT homotypic NF-κB-genetic variant viral strains.

### Generation of pseudotyped reporter viral stocks

Pseudotyped viral stocks were produced by co-transfecting each variant viral construct and third-generation lentiviral packaging vectors in HEK293T cells using a standard calcium phosphate protocol (42) (PMID: 8604299). HEK293T cells at 40% confluency seeded in a 6-well culture plate were transfected with a plasmid DNA cocktail consisting of 3 μg of the viral vector, 1.5 μg of psPAX2 (Catalog no: 11348; NIH AIDS reagent program, USA), 0.9 μg of VSV-G (Catalog no: 4693; NIH AIDS reagent program, USA), and pREV-1 (Catalog no. 1443; NIH AIDS reagent program, USA) along with 0.05 μg of pmScarlet_C1 (Catalog no. 85042; Addgene), the latter as the transfection control. Six hours following transfection, the medium was replenished with the complete DMEM medium. Forty-eight hours following transfection, the culture supernatant containing viral particles was harvested. The supernatant was passed through a 0.22 μm filter and stored at −80°C for future use.

### Estimation of the Relative Infectious Units (RIU) of viral stocks

Approximately 1 ✕ 10^6^ Jurkat T-cells per well seeded in a 12-well culture plate were infected with viral stocks serially diluted in a 5-fold series (from 5X to 625X) in a total volume of 1 ml of complete RPMI medium supplemented with 25 μg/ml of DEAE-Dextran (Catalog no. 30461; Sigma Aldrich). Six hours following the infection, the cells were washed and resuspended in 1 ml of complete RPMI medium and incubated for 48 h. The cells were activated with a cocktail of T-cell activators consisting of 40 ng/ml tumor necrosis factor-alpha (TNF-α; Catalog no. T0157; Sigma-Aldrich), 40 ng/ml phorbol myristate acetate (PMA; Catalog no. P8139; Sigma-Aldrich), and 2.5 mM hexamethylene bis-acetamide (HMBA; Catalog no. 224235; Sigma-Aldrich) for 24 h and analyzed for fluorescence expression using a flow cytometer (BD FAC AriaIII sorter; BD biosciences, New Jersey, USA). A viral titration curve was constructed using viral dilutions on the X-axis and percentage EGFP^+ve^ cells on the Y-axis. Using regression analysis, we identified the viral titer necessary to cause 5-10% of EGFP^+ve^ cells that would correspond to 0.05-0.1 RIU.

### Generation of stable polyclonal populations and clonal cell lines

#### (i) Stable-cell pools of cLdGIT LTR-variant viral strains/ NF-κB genetic-variant viral strains

Jurkat T-cells were infected with viral stocks producing approximately 10% infectivity and incubated for 48 h. Following infection, the cells were activated using a cocktail of T-cell activators for 24 h, and EGFP^+ve^ cells were sorted and expanded. The purity of sorted cells was validated using a small aliquot of the sorted cells. The sorted and expanded cells are expected to represent a mixed population with random proviral integration sites.

#### (ii) Clonal cell lines of cLdGIT LTR-variant panel

Jurkat T-cells were infected with individual viral strains of the cLdGIT panel and activated for 24 h using a cocktail of T-cell activators. Following activation, single cells expressing fluorescence were sorted into individual wells of a 96-cluster plate containing 100 μl of complete RPMI medium supplemented with 50% spent medium. Two weeks following sorting, the cells were transferred to 48-well culture plates and subsequently to 24 well-culture plates. The cells were allowed to expand for four weeks and stored or used in experiments.

#### (iii) Stable-cell pools of Tet-ON cLdG NF-κB genetic-variant viral strains

The Tet-ON Tat cell line expressing Tat under the control of Tet-ON promoter was generated in two successive steps. First, a stable cell line expressing rtTA3 was generated using puromycin selection (43) (PMID: 33101270). This was followed by infection of the stable rtTA3 expressing cell line with Tet-ON Tat vector where Tat expression was placed under the control of Tet-ON promoter (Saini C et al., unpublished) to generate a clonal cell line. The Tet-ON Tat expressing cell line was infected with the individual cLdG NF-κB variant viral strains of the panel at <0.1 RIU for 48 h. Following infection, the cells were activated with 800 ng/ml Doxycycline (Dox) and a cocktail of T-cell activators for 24 h. The cells expressing EGFP were sorted and expanded for further experiments. A small aliquot of the sorted cell pools was analyzed by flow cytometry to confirm the purity of the sorted cells.

### Measurement of gene expression noise

Gene expression noise of a stable-cell pool was measured by estimating the proportion of cells present in the ‘Mid’ region, the region with intermediate/low level of gene expression between the OFF and High gene expression states that exhibit the maximum effect of stochasticity. A gating strategy was optimized by overlaying the histogram profiles of non-infected (open histogram) and the infected (solid histogram) cell populations to identify cells under the ‘OFF’, ‘ON’, ‘Mid’, and ‘High’ regions. The relative fluorescence unit (RFU) of the OFF-region encompassed >95% of the total cells in the non-infected cell population (~10-300 RFU), and the ON-region marker spanned the total EGFP^+ve^ cells (~300-200,000 RFU) in the infected cell population. The High-region marker spanned >95% of the cells in the ON-region of the infected cell population (~5,000-200,000 RFU). The intermediate region between OFF- and High-regions was considered the Mid-region (~300-5,000 RFU). Gene expression noise was monitored by measuring the Mid:ON ratio, the proportion of cells in the Mid-region relative to the cells present in the ON-region (32) (PMID: 19132086). Gene expression noise of the clonal population was measured by estimating the heterogeneity of EGFP expression in the population. We used the coefficient of variance (CV), the ratio of standard deviation to the mean of EGFP expression, to measure the heterogeneity of EGFP expression in the population (44) (PMID: 16179466).

### Inhibition of the NF-κB pathway and Western blot analysis

Stable-cell pools infected with individual LTR-variant viral strains of a panel were treated with a cocktail of T-cell activators with/without 1 μM of MG-132 (Catalog no. M7449; Sigma-Aldrich), an inhibitor of the NF-κB pathway for 12 h. Subsequently, the cells were stained for dead cell exclusion, and the fluorescence was analyzed using a flow cytometer.

To evaluate the inhibitory effect of MG-132 on the NF-κB pathway, Jurkat T-cells were pre-treated with/without MG-132 for 12 h and subsequently treated with a cocktail of T-cell activators. Whole-cell lysates were prepared at 0, 15, 30, 60 min following activation using RIPA lysis buffer consisting of 50 mM Tris–HCl, pH 7.4 (Tris-Catalog no. 15965; Fisher Scientific, India; HCl-Catalog no. HC301585; Merck, India), 150 mM NaCl (Catalog no. S6191; Sigma, India), 0.1% SDS (Catalog no. L3771; Sigma-Aldrich, India), 1% Triton X-100 (Catalog no. 845; HiMedia, India), 0.25% sodium deoxycholate (Catalog no. D6750-100G; Sigma, India), and a protease inhibitor cocktail (Catalog no. 786-437; G-Biosciences) for 30 min. The lysate was centrifuged at 13,000 rpm for 5 min, and the supernatants were collected. Protein concentration was estimated by the Bradford assay using a commercial kit (Catalog no. 5000001; Bio-Rad, India). For the Western blot analysis, the cell lysate was electrophoresed using SDS-PAGE (5% stacking and 12% resolving gel) for ~2 h. The separated proteins were transferred to a 0.2 μm PVDF membrane (Catalog no. BSP0161; Pall Corporation) for 1 h and 20 min at 90 V and 4°C. Following the transfer, the membrane was blocked with 5% skimmed milk (Catalog no. GRM1254; HiMedia) at room temperature for 1 h and washed once with 1X phosphate-buffered saline (PBS). Proteins were probed with an anti-IκBα antibody raised in rabbits (Catalog no. ab32518; Abcam) and anti-GAPDH antibodies raised in mice (Catalog no. 10-10011; Abgenex). After overnight incubation, the blots were washed three times for 10 minutes each time, using 1X PBS supplemented with 0.5% Tween-20 (PBST) (Catalog no. TC287; HiMedia, India). The membrane was incubated with Horseradish peroxidase-conjugated (HRP-conjugated) anti-rabbit (Catalog no. 1721019; Bio-Rad, India) or anti-mouse (Catalog no. 1706516; Bio-Rad, India) antibodies in 1% skimmed milk for 1 h at room temperature. Proteins were detected using Enhanced-Luminol-based chemiluminescent substrate (ECL; Catalog no. 1705060; Bio-Rad, India) and visualized using an I-Bright CL1000 Chemidoc system (Thermofisher). Blots were quantified using ImageJ software (v1.52a; NIH, U.S.A).

### Analysis of Tat and p65 transcripts

Tat and p65 transcripts were quantitated using real-time qPCR. GAPDH transcript levels were analyzed as the housekeeping control. Total RNA was extracted from 1 ✕ 10^6^ cells using the TRI reagent (Catalog no. T9424; Sigma-Aldrich). From the extracted RNA, 2 μg RNA was treated with DNase (Catalog no. EN0525; Thermo Scientific) for 30 min at 37°C to remove genomic DNA contamination. The DNAse was inactivated by incubating the sample at 65°C for 10 min. Approximately, 500 ng of DNase-treated RNA was converted into cDNA using Oligo-dT primers in a reaction volume of 10 μl using the OneScript Plus cDNA synthesis kit (Catalog no. G236; ABM). The cDNA was amplified using primer pairs specific to Tat, p65, or GAPDH transcripts (the primer details are presented in Table 3). The PCR was performed using the BioRad CFX96 Real-Time PCR cycler using a DyNAmo Flash SYBR green qRT-PCR kit (Catalog no. F415XL; Thermo Scientific).

**Table 3.**
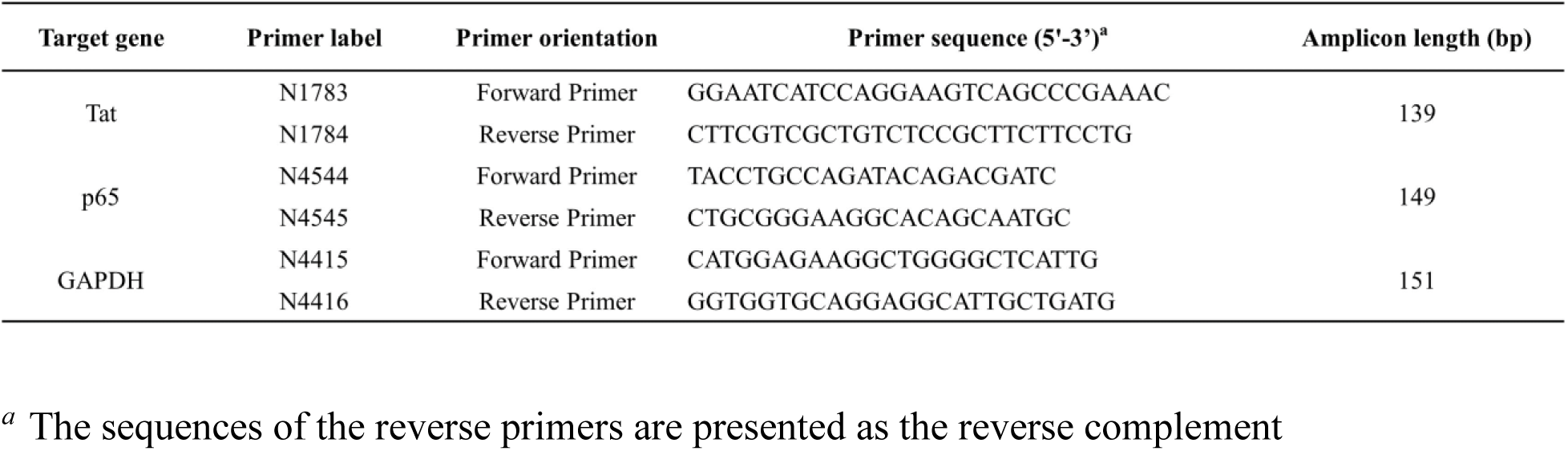
Primer sets used for transcript analysis using qRT-PCR.

### Kinetic viral gene expression analysis

The stable-cell pool of cLdGIT HHC was activated with a cocktail of T-cell activators in a standard homogeneous environment to demonstrate the unstable nature of the cells in the Mid region. An aliquot of cells was collected every four hours, and the cells were stained for dead cell exclusion and analyzed by flow cytometry for EGFP expression for 24 h. Kinetic profiles of the cell percentage in the OFF-, Mid-, and High-regions were constructed. The latent population was sorted from the stable-cell pools of cLdGIT LTR variant viral strains to generate kinetic curves of EGFP expression. Following sorting, the cells were homogeneously activated using a cocktail of T-cell activators, and a small aliquot of the cells was analyzed using flow cytometry every 2 h for 22 h. Kinetic profiles of median fluorescence intensity (MFI) of EGFP^+ve^ cells were plotted.

### Cell viability assay

Approximately 1 ✕ 10^6^ cells were harvested and washed once with 1X PBS. The cells were suspended in 100 μl of 1:10,000 of the Live/Dead stain (Catalog no. L34974; Invitrogen) and incubated in the dark for 30 min at 4°C. Following incubation, the cells were washed with 1X PBS supplemented with 2% FBS and resuspended in 200 μl of 1X PBS containing 2% FBS. The cells were analyzed using a flow cytometer.

### Data analysis, plotting, and statistical evaluation

All flow cytometry data analyses were performed using FCS Express 6 software (De Novo Software, Los Angeles, CA). The data for all the experiments were plotted using GraphPad Prism software (Version 5.02 and Version 9.3.1). The Hill coefficient (H) value was derived by fitting the MFI kinetic trajectory from flow cytometry analysis in a Hill equation curve using GraphPad Prism software. Statistical evaluation for all the experiments was performed using the same software. The statistical tests performed for each experiment are depicted in the corresponding figure legends, along with the *p*-values.

## Results

### The infected cells in the ‘Mid’ region are unstable

HIV-1 promoter displays a relatively higher level of gene expression noise than eukaryotic promoters (21,22) (PMID: 23064634, 20409455). Experimental and simulation studies using HIV-1 viral promoter demonstrated that infected cells with low/intermediate gene expression levels exhibit maximum stochasticity effect (14) (PMID: 16051143). The variations in the TFBS profile of HIV-1C LTR offer a unique advantage in asking how such changes could modulate transcriptional or gene expression noise of the viral promoter when the extrinsic factors are kept constant. For example, the variation in the number of NF-κB binding sites in HIV-1C, but not in other HIV-1 subtypes, offers an opportunity to examine the association between the number of NF-κB motifs in the enhancer region of the LTR and gene expression noise. Of note, we previously demonstrated a positive correlation between the number of NF-κB motifs varying between 1 to 4 and the transcriptional activity of the LTR (34) (PMID: 23132857).

To this end, we used the cLdGIT vector; a minimal sub-genomic reporter vector described previously (39) (PMID: 32669338). The cLdGIT viral vector contains both the LTR and Tat of HIV-1C origin. The LTR, being representative of the subtype, contains three NF-κB motifs in the enhancer region and regulates the co-expression of a d2EGFP and the Tat protein under an IRES element (Fig 1A). Minimal HIV-1 viral vectors encoding reporter genes have been widely used to study HIV-1 latency (14,22) (PMID: 16051143, 20409455). Viral stocks pseudotyped with VSV-G envelope were produced in HEK293T cells. Jurkat T-cells were infected at a low multiplicity of infection (MOI>0.1), yielding a polyclonal population with a broad range of single viral integration per cell (Fig 1B). The infected cells were activated for 24 h with a cocktail of T-cell activators consisting of 40 ng/ml PMA, 40 ng/ml of TNF-α, and 2.5 mM HMBA. The cells were analyzed by flow cytometry. A gating strategy was optimized by overlaying the histogram profiles of non-infected and infected cell populations to identify cells under the ON and OFF states (Fig 1C). Since cells from the Mid region are subjected to maximum stochastic fluctuations (14) (PMID: 16051143), we sorted cells from this region and monitored the EGFP fluorescence every four hours up to 24 hours, using flow cytometry.

**Fig 1:**
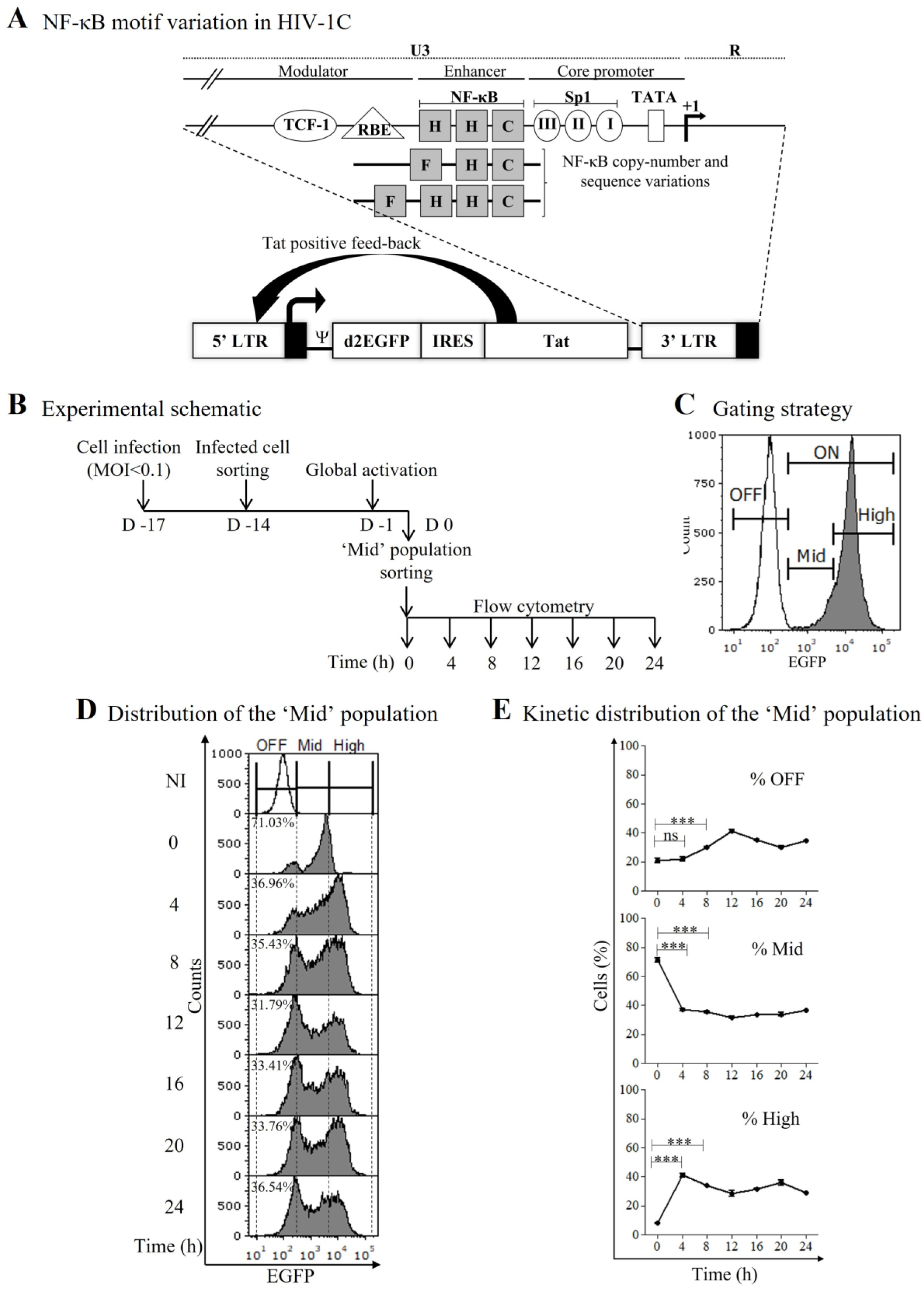
The stochastic nature of the ‘Mid’ population in HIV-1C infection. (A) Schematic representation of NF-κB motif variation in HIV-1C LTR and the sub-genomic viral construct used in the present study. The top panel represents a simplified version of the U3 regulatory region of the LTR. Several important TFBS located in the modulator, enhancer, and core promoter regions have been labeled. C, F, and H represent the three genetically distinct NF-κB motifs (34). While the canonical HIV-1C LTR contains three NF-κB binding motifs (HHC), strains containing three copies of genetically distinct NF-κB motifs (FHC) and four copies of the TFBS (FHHC) are also common. A viral strain containing three canonical NF-κB sites was constructed using the LGIT vector, a bi-cistronic sub-genomic reporter viral construct. In these vectors, both Tat and the LTR are of HIV-1C origin. (B) Experimental schematic. Jurkat T-cells were infected with the cLdGIT HHC virus at an MOI of <0.1, and 72 h later the cells were sorted using a flow cytometer. Two weeks following relaxation and expansion, the cells were induced for 24 h with a cocktail of T-cell activators (TNF-α, PMA, HMBA), and the intermediate ‘Mid’ population was sorted. Following the sorting, EGFP expression was monitored every 4 h for 24 h using a flow cytometer. (C) The gating strategy to demarcate the ‘OFF’, ‘ON’, ‘Mid’, and ‘High’ regions. The histogram represents an overlaid gene expression profile of the non-infected (open histogram) and infected (solid histogram) cell populations. The intermediate region between ‘OFF’ and ‘High’ is considered the ‘Mid’ region (~300-5,000 RFU). These RFU values were set based on profiles observed in multiple replicate experiments. (D) Representative stacked histogram profile of the sorted ‘Mid’ population across time. The dotted line delineates the OFF, Mid, and High regions. The percentages of the Mid-region cells across time are depicted in the histogram plots. NI, no infection. (E) Line graphs depict the distribution of the total sorted cells from the ‘Mid’ region into the ‘OFF’, ‘Mid’, and ‘High’ regions. Data are representative of three independent experiments. Mean values from experimental triplicates ± standard deviation (SD) are plotted. One-way ANOVA followed by Bonferroni’s multiple comparison test was used for statistical evaluation (***, p<0.0001; ns, not significant).

The sorted cells from the Mid region were highly unstable and transitioned to the OFF and High regions (Figs 1D and 1E), consistent with the previous report (14) (PMID: 16051143). For example, the percentage of cells in the Mid region of fluorescence dropped from 71.03 ± 1.45% to 31.79 ± 0.97% between 0 and 12 h, respectively. In contrast, the percentages of cells in the OFF region increased from 20.93 ± 1.32% to 41.16 ± 1.10%, and those in the High region also increased from 8.00 ± 0.25% to 28.71 ± 1.97% at the same time points, respectively. Thus, the cells of the Mid region transitioned to the other two regions conforming to the highly unstable nature of the region.

### Gene expression noise is inversely correlated to the number of NF-κB motifs in the LTR

The present study attempts to understand the effect of varying transcriptional strength of HIV-1 LTR on gene expression noise. HIV-1C contains three canonical NF-κB binding sites in the enhancer region of the promoter, a number higher than in other HIV-1 subtypes. We also demonstrated the frequent occurrence of 4-κB viral strains in the population (34) (PMID: 23132857). Further, a direct correlation exists between the number of NF-κB binding sites and the transcriptional strength of the viral promoter. Hence, the modulation of gene expression noise could be evaluated by varying the number of the κB-motifs that, in turn, alter the transcriptional strength of the LTR. To this end, we generated a panel of cLdGIT viral vectors with a varying copy number of NF-κB binding motifs ranging from 0 to 4 by introducing inactivating point mutations into individual binding motifs (Fig 2A). Jurkat T-cells were infected at an MOI of <0.1 and activated with a cocktail of T-cell activators for 24 h (Fig 2B). Cells were stained for dead-cell exclusion, and fluorescence expression was analyzed by flow cytometry.

**Fig 2:**
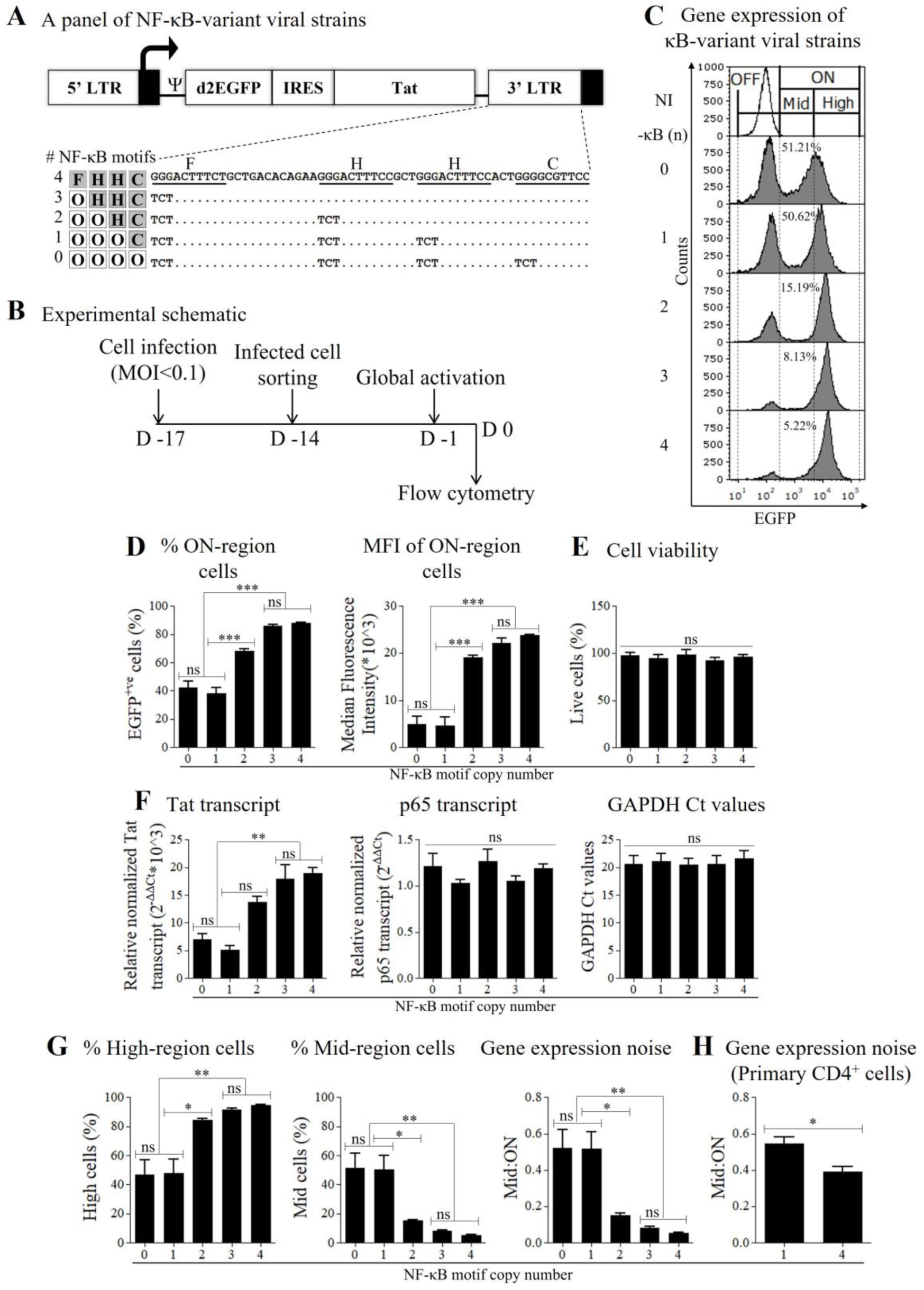
An inverse correlation between the number of NF-κB motifs and gene expression noise of HIV-1C LTR. (A) A panel of cLdGIT sub-genomic variant viral strains containing a varying number of NF-κB motifs in the HIV-1C LTR. The nucleotide sequences of the NF-κB motifs are underlined, and the inactivating point mutations introduced are shown. The dots represent sequence identity. (B) Experimental schematic. Following infection and sorting, infected Jurkat T-cells were activated with a cocktail of T-cell activators (TNF-α, PMA, HMBA) for 24 h and analyzed for dead-cell exclusion and EGFP expression using flow cytometry, and Tat and p65 transcripts by qRT-PCR. (C) Representative stacked histogram profiles of EGFP expression of the variant viral strains at 24 h following activation. The dotted line delineates the OFF, Mid, High, and ON regions. The open histogram represents uninfected and activated cells, and solid histograms represent infected and activated cells. The percentages of the Mid-region cells corresponding to each LTR-variant viral strain are depicted in the histogram plots. NI, no infection. (D) Percentage (left panel) and MFI (right panel) values of the total EGFP^+ve^ cells present in the ON region and (E) the percentage values of activated live cells relative to their no activation control from three independent experiments ± standard error of the mean (SEM) are plotted. One-way ANOVA followed by Bonferroni’s multiple comparison test was used for statistical evaluation (***, p<0.0001; ns, not significant). (F) The relative normalized mean values of Tat, p65 mRNA transcripts, and GAPDH Ct values are plotted. Data represent mean values from two independent experiments ± SEM. One-way ANOVA followed by Bonferroni’s multiple comparison test was used for statistical evaluation (**, p<0.001; ns, not significant). (G) The percentage values of EGFP^+ve^ cells in the High (left panel) and Mid (middle panel) regions and gene expression noise measured using the Mid:ON ratio (right panel) of the variant viral strains are plotted. Data represent mean values from three independent experiments ± SEM. One-way ANOVA followed by Bonferroni multiple comparison test was used for statistical evaluation (**, p<0.001; *, p<0.05; ns, not significant). (H) Gene expression noise analysis in primary CD4^+^ T-cells, derived from three healthy subjects using 1- and 4-κB variant viral strains. Mean Mid:ON ratio values from three independent experiments performed on three subjects ± SEM are plotted. A two-tailed unpaired Student’s *t*-test was used for statistical evaluation (*, p<0.05).

The stacked histogram profile represents the activated populations of non-infected (open histogram) and infected (solid histogram) Jurkat T-cells, corresponding to all the five LTR-variant viral strains (Fig 2C). The cell population of each variant viral strain was spread spanning the OFF, Mid, and High regions depending on the intensity of EGFP expression. The percentages of cells in the ON region, i.e., the total EGFP^+ve^ cells, were measured using the stacked histogram profile (Fig 2D, left panel). A proportional increase in the percentage of EGFP^+ve^ cells was evident with the increasing number of NF-κB motifs in the LTR, although the percentages of the infected cells following the sorting were comparable among the variant viral strains. The LTRs containing three and four NF-κB motifs demonstrated the highest percentages of EGFP^+ve^ cells, 85.92 ± 0.71 and 87.74 ± 0.55, respectively, the difference between the two LTRs being insignificant. In contrast, the 0-κB LTR (in which all four κB-motifs have been inactivated) and 1-κB LTR demonstrated the lowest percentages of EGFP^+ve^ cells, 42.22 ± 3.64 and 37.97 ± 3.34, respectively. In contrast, the 2-κB LTR with 67.92 ± 1.41% EGFP^+ve^ cells occupied an intermediate position. Likewise, the MFI of the ‘ON’ population demonstrated a similar trend (Fig 2D, right panel). The 3-κB and 4-κB LTRs showed the highest relative fluorescence, 22,143.00 ± 890.90 RFU, and 23,772.23 ± 153.12 RFU, respectively, compared to the 0-κB and 1-κB promoters, 4,871.79 ± 1,333.21 RFU and 4,526.37 ± 1,565.78 RFU, respectively. At the same time, the 2-κB LTR showed an intensity of intermediary level, the corresponding MFI value being 19,065.01 ± 340.77 RFU. Of note, the MFI value, but not the percentage, of the 2-κB LTR did not differ significantly from those of 3- and 4-κB LTRs. Notably, the percentage cell viability was comparable among all the κB-variant viral strains (Fig 2E).

Like the percentages of EGFP^+ve^ cells and MFI profiles, the Tat transcript levels also increased proportionately with the increasing number of the κB-motifs in the LTR (Fig 2F, left panel). Additionally, as expected, the intracellular p65 transcript levels were constant among the variant viral strains (Fig 2F, middle panel). GAPDH transcript levels were used as an internal cellular control to validate Tat and p65 transcript expression (Fig 2F, right panel). Thus, the difference in transcriptional strength between strong (2-, 3-, and 4-κB) and weak (0-, 1) LTRs could be attributed to the difference in NF-κB motif copy number when all other external factors remain constant.

Extending the analysis further, we evaluated how the κB-motif copy number variation in the LTR would modulate the percentages of High- and Mid-region cell populations within the ON region since a ratio of these two compartments would represent gene expression noise. The data showed that the EGFP^+ve^ percentages of the High-region cells (Fig 2G, left panel) were directly proportional to the number of the κB-motifs in the LTR. For example, 46.92 ± 8.27%, 47.59 ± 7.90%, 84.23 ± 1.06%, 91.50 ± 0.69%, and 94.63 ± 0.46% High-region cells, were EGFP^+ve^ corresponding to 0-, 1-, 2-, 3-, and 4-κB LTRs, respectively. This result reflects that the stronger the LTR, the higher Tat production, and hence the larger the transactivated population. In contrast, the profile of EGFP^+ve^ cells in the Mid region was inversely correlated to the number of the κB-motifs in the LTR. Thus, 51.21 ± 8.32%, 50.62 ± 7.70%, 15.19 ± 0.67%, 8.13 ± 0.60%, and 5.22 ± 0.37% Mid-region cells, were EGFP^+ve^ corresponding to 0-, 1-, 2-, 3-, and 4-κB LTRs, respectively (Fig 2G, middle panel). An increase in the proportion of cells in the Mid region cannot be attributed to autofluorescence of the dead cells since the cell populations were stained for dead-cell exclusion.

Our data are consistent with the previous reports, which demonstrated that the Mid population with intermediate levels of gene expression was highly unstable, contributing to the High or OFF states (14,32,33) (PMID: 16051143, 19132086, 23874178). Considering the Mid:ON ratio as a metric to measure the level of randomness in gene expression (32) (PMID: 19132086), we compared the profile of gene expression noise with the varying number of NF-κB binding motifs. Inactivation of NF-κB motifs is expected to increase the Mid:ON ratio. Consistent with the expectation, we found an inverse correlation between the number of NF-κB motifs and gene expression noise (Fig 2G, right panel). The Mid:ON ratio decreased significantly as the number of κB-motifs increased from 0 to 4 in the LTR. For example, the noise levels of 0-, 1-, 2-, 3-, and 4-κB LTRs were 0.52 ± 0.08, 0.52 ± 0.07, 0.15 ± 0.01, 0.08 ± 0.01, and 0.05 ± 0.004, respectively. Notably, the noise levels between the 3- and 4-κB variant LTRs were comparable. In summary, our data show that gene expression noise of HIV-1 LTR decreases with increased transcriptional strength in Jurkat T-cells. We observed a similar trend of noise difference between 1- and 4-κB variant viral strains in primary CD4^+^ T-cells, although the difference was modestly significant (Fig 2H). Of note, the two natural and canonical HIV-1C LTRs containing 3- or 4-κB motifs did not differ significantly in any profile, including in gene expression noise. For practical reasons, we will henceforth refer to the 0- and 1-κB LTRs as weak promoters and 3- and 4-κB LTRs as strong promoters. The 2-κB LTR largely behaved as a strong LTR, although this LTR mostly occupied an intermediate position in many properties.

### Gene expression noise variability is consistent among clonal cell populations

The analysis of gene expression noise performed above using infected cell pools offers the advantage of normalizing the integration site differences. Genome integration sites can influence viral gene expression (45) (PMID: 20941390). Evaluating gene expression noise using cell pools may mask minor or moderate variations due to genome integration site differences. We, therefore, examined the profiles of gene expression noise modulation of the clonal population of cells infected with κB-motif number variant viral strains. Individual EGFP^+ve^ cells were sorted from the ON region of the cLdGIT LTR-variant viral strains into a 96-well culture plate, and the clones were expanded for ~4 weeks (Fig 3A). The clonal populations (30, 14, 26, 34, and 49 clones representing 0-, 1-, 2-, 3-, and 4-κB variant viral strains, respectively) were treated with a cocktail of T-cell activators for 24 h that induced the proviruses to the highest magnitude of activation.

**Fig 3:**
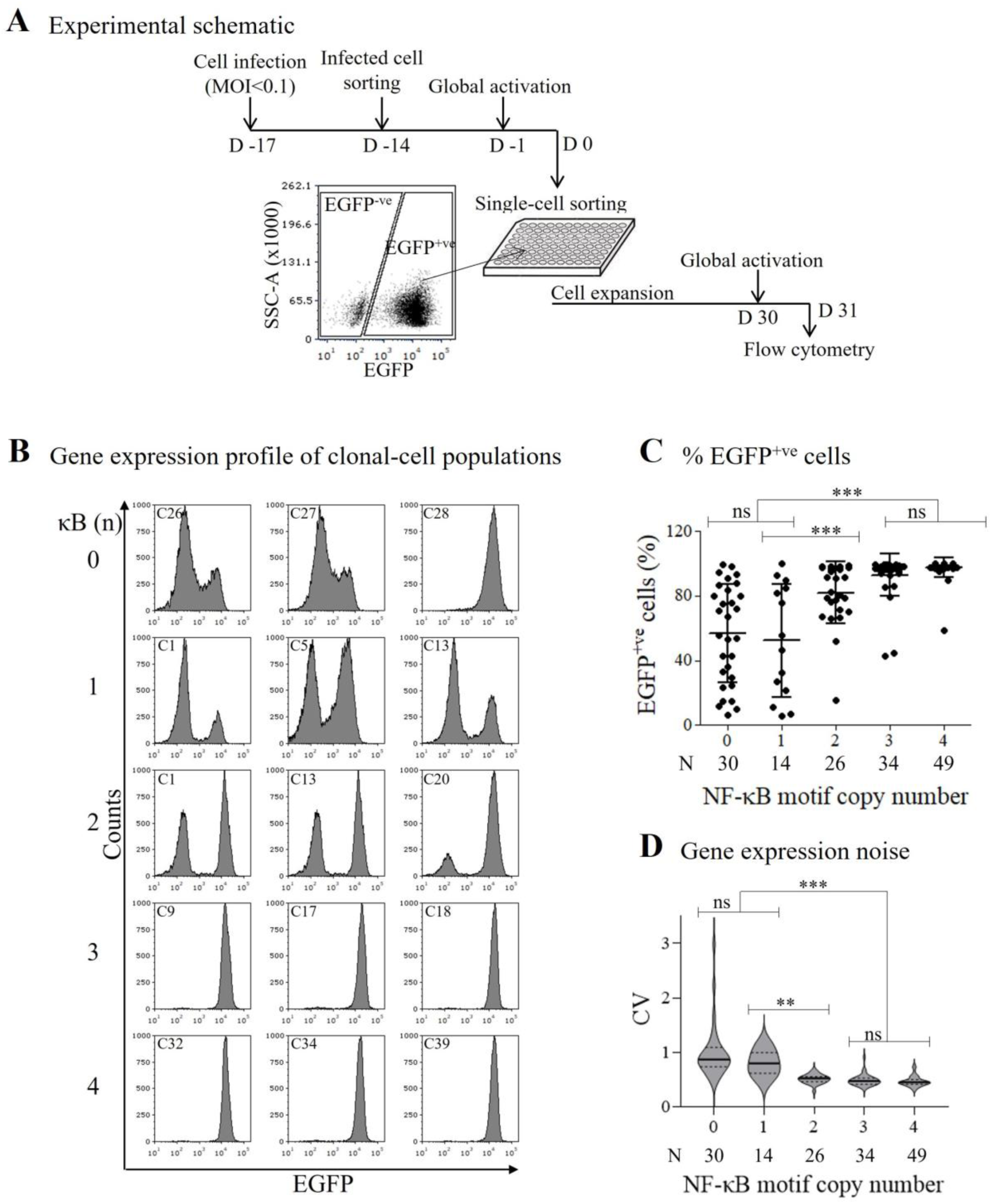
The profile of gene expression noise of clonal cell populations of κB-variant strains. (A) Experimental schematic of gene expression noise analysis in clonal cell populations. The infected polyclonal cell pool was activated for 24 h, and single, EGFP-expressing cells were sorted into a 96-well culture plate using a flow sorter. The cells were expanded for ~4 weeks, activated with a cocktail of T-cell activators for 24 h, and analyzed using a flow cytometer. (B) Gene expression profile. Histogram profiles of three representative clonal-cell populations for each LTR-variant viral strain. (C) The percentage EGFP^+ve^ cell profiles of κB-variant viral strains. Each dot represents the mean percentage of EGFP^+ve^ cells from experimental triplicates of a clonal population. One-way ANOVA with Bonferroni’s multiple comparison test was performed for statistical evaluation (***, p<0.0001; ns, not significant). N, number of clones analyzed per variant viral strain. (D) Violin plots depicting gene expression noise measured as coefficient of variance (CV) of clonal populations of each variant viral strain. Mean CV values using experimental triplicates of each clonal population for all the variant viral strains are plotted. Welch’s ANOVA test with Dunnett’s T3 multiple comparisons test was performed for statistical evaluation (***, p<0.0001; **, p<0.001; ns, not significant). N, number of clones analyzed per variant viral strain.

Following cell activation, most of the clonal cell populations expressed EGFP. The gene expression profiles of a few representative clonal populations of each LTR-variant viral strain are presented (Fig 3B). However, significant differences were evident in the percentages of EGFP^+ve^ cells depending on the NF-κB motif copy number (Fig 3C). The mean percentage values of cells expressing fluorescence in clonal populations significantly differed between the weak (0- and 1-κB) and strong (3- and 4-κB) LTRs. Importantly, for 0- and 1-κB LTRs, the variation in the percentages of EGFP^+ve^ cells expanded over a broad range, and for the other three LTRs, the spread was the narrowest. For example, the mean percentages of EGFP^+ve^ cells of clonal populations of 0- and 4-κB LTRs were 56.95 ± 30.53, and 97.49 ± 5.90, respectively, with the difference being significant. Thus, a larger percentage of cells representing the strong LTRs were EGFP^+ve^ than those of the weak LTRs.

To appreciate the association between the transcriptional strength of the LTR and gene expression noise, we estimated the CV of each clonal population. CV, defined as the ratio of standard deviation to the population mean, normalizes variability across mean values, thereby enabling the comparison of variability among clonal populations (44) (PMID: 16179466). Since heterogeneity in gene expression in a clonal cell population is a consequence of gene expression noise, we estimated the variation in gene expression in the clonal population. As is evident from the analysis, the mean CV values of the clonal populations of the two weak LTRs were the highest, with 0- and 1-κB LTRs showing 1.00 ± 0.51, 0.81 ± 0.23, respectively. In contrast, the three other LTRs (2-, 3-, and 4-κB LTRs) demonstrated mean CV values 0.51 ± 0.08, 0.49 ± 0.10, and 0.47 ± 0.10, respectively (Fig 3D). From these data, it is evident that the presence of more copies of the NF-κB motif augments the transcriptional strength of the LTR proportionately and reduces variability in gene expression or gene expression noise correspondingly. The two strong LTRs containing 3- or 4-κB motifs circulating in the population demonstrate the highest transcriptional activity, indistinguishable between the two. Thus, at least in HIV-1C, the stronger transcriptional activity of the LTR, characterized by the lowest gene expression noise, appears to be associated with a replication advantage. Notably, the near uniformly high-level gene expression of the strong LTRs implies that the integration site may not be a major factor, at least in selected clonal populations, contributing to the gene expression noise of strong viral promoters. In summary, the presence of more copies of the NF-κB motif, on the one hand, appears to augment the transcriptional strength of the LTR and, on the other hand, reduce gene expression noise. These qualities may modulate HIV-1 latency considerably.

### Attenuating the NF-κB signaling pathway augments gene expression noise of the strong LTRs

Our data demonstrated a negative correlation between the copy-number of NF-κB motifs in the LTR and gene expression noise (Fig 2G, right panel). Stronger LTRs containing 3- or 4-NF-κB binding sites demonstrated the lowest magnitude of gene expression noise (Fig 3D). We next attempted to understand whether the depletion of the nuclear NF-κB concentration could modulate gene expression noise of strong LTRs. To this end, we used MG-132, a small-peptide inhibitor of IκBα degradation, thereby preventing the translocation of NF-κB from the cytoplasm to the nucleus (46) (PMID: 21627972). Progressively increasing concentrations of MG-132 are expected to reduce the intracellular NF-κB levels concomitantly, thus augmenting gene expression noise of the LTR.

Jurkat T-cells were treated with MG-132, at concentrations ranging from 0.25 – 2.0 µM for 12 h, and cell viability was evaluated by flow cytometry (Fig 4A). We next examined the magnitude of inhibition of the NF-κB signaling pathway at different concentrations of MG-132. Jurkat T-cells pre-treated with different concentrations of MG-132 for 12 h were induced with a T-cell activator cocktail. The cells were harvested for Western blot analysis at 0, 15, 30, and 60 min and were probed for the IκBα protein. We found that pre-treatment of MG-132 at a final concentration of 1.0 µM, but not below, resulted in significant inhibition of NF-κB signaling (Fig 4B). Cells with or without activation or MG-132 treatment were also included for comparison. In the absence of cell activation, the intracellular IκBα levels remained comparable without considerable variation (Fig 4B, left panel). Jurkat T-cells activated but not treated with MG-132 showed depletion of intracellular IκBα levels within 15 min, which reduced further at 30 min and started to recover by 60 min (Fig 4B, middle panel), suggesting efficient activation of the NF-κB pathway. Notably, the activation of the NF-κB pathway was abrogated when the cells were pre-treated with MG-132, as a reduction in the IκBα levels was not seen at any time points tested (Fig 4B, right panel).

**Fig 4:**
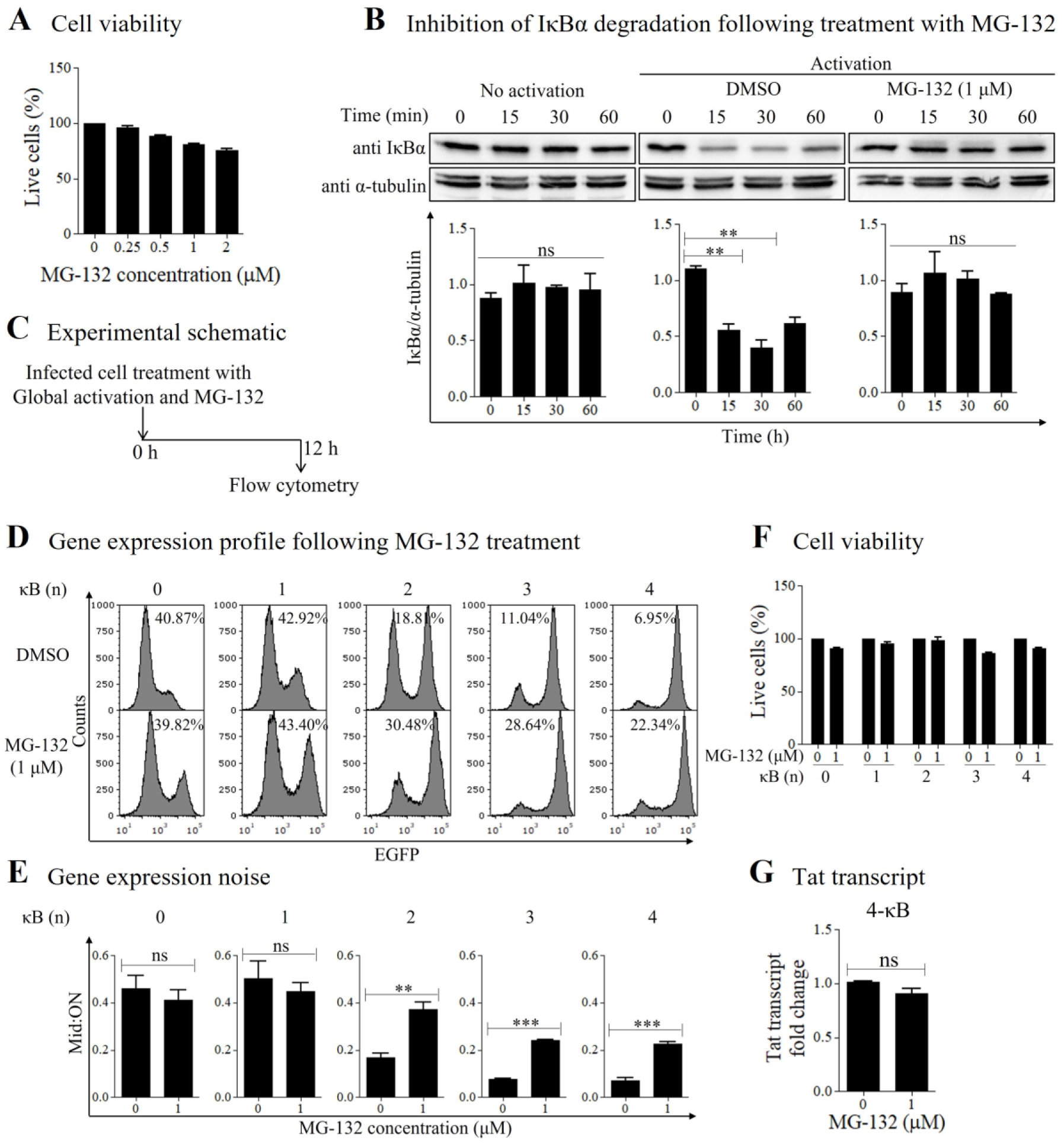
Augmented gene expression noise of strong LTRs following the inhibition of the NF-κB signaling pathway. (A) Cell viability following MG-132 treatment. Jurkat T-cells were treated with a range of MG-132 concentrations for 12 h, followed by staining for dead-cell exclusion and flow cytometry analysis. Percentages of live cells are plotted for the range of MG-132 concentrations. Data represent mean values from three independent experiments ± SEM. (B) Representative Western blot images demonstrating the kinetics of IκBα expression in Jurkat T-cells following no treatment/treatment with a cocktail of T-cell activators and/or MG-132. α-tubulin expression was used as the loading control. The bottom panel demonstrates the mean densitometric analysis of Western blot images from two independent experiments ± SEM. One-way ANOVA with Bonferroni’s multiple comparison test was performed for statistical evaluation (**, p<0.001; ns, not significant). (C) Experimental schematic for gene expression noise analysis. Infected cells were treated with MG-132 and a cocktail of T-cell activators (TNF-α, PMA, HMBA) for 12 h and analyzed using flow cytometry. (D) Representative stacked histogram profiles of EGFP expression of the panel of κB-variant viral strains following treatment with DMSO (top panel) or MG-132 (bottom panel). The percentages of the Mid-region cells corresponding to each LTR-variant viral strain with/without MG-132 treatment are depicted in the histogram plots. (E) Gene expression noise measured as Mid:ON ratio for the respective variant viral strains. Mean Mid:ON ratio values from three independent experiments ± SEM are plotted for both DMSO- and inhibitor-treated cells for each variant viral strain. A two-tailed unpaired Student’s *t*-test was performed for statistical evaluation (***, p<0.0001; **, p<0.001; ns, not significant). (F) Percentage cell viability of all the LTR-variant viral strains following treatment with a cocktail of global activators, with/without treatment with MG-132. Mean values from three independent experiments ± SEM are plotted. (G) Tat transcript analysis of 4-κB LTR following treatment with a cocktail of global activators, with/without treatment of MG-132. Mean values from three independent experiments ± SEM are plotted. A two-tailed unpaired Student’s *t*-test was performed for statistical evaluation (ns, not significant).

Using the optimized experimental conditions described above, we evaluated the effect of MG-132 on gene expression noise of variant LTRs. Jurkat T-cells, individually infected with LTR-variant viral strains, were treated with MG-132 or DMSO, induced with a cocktail of T-cell activators for 12 h, and analyzed using flow cytometry (Fig 4C). The analysis showed that NF-κB inhibition profoundly impacted gene expression noise of the strong LTR variant viral strains. Twelve hours following activation and without MG-132 treatment, cells of strong LTRs (3- and 4-κB), but not weak LTRS (0- and 1-κB), transited out of the Mid region rapidly (Fig 4D, top panel). Importantly, when NF-κB signaling was blocked using MG-132, cell migration of the three strong LTRs (2-, 3-, and 4-κB), but not the two weak LTRs (0- and 1-κB), reduced significantly out of the Mid region (Fig 4D, bottom panel). For example, while in the absence of MG-132, 18.81 ± 1.24%, 11.04 ± 1.41%, and 6.95 ± 0.88% cells were present in the Mid region of 2-, 3-, and 4-κB LTRs, respectively; in the presence of the inhibitor, these percentages increased to 30.48 ± 5.06%, 28.64 ± 5.60%, and 22.34 ± 0.75%, respectively. The increased percentage of cells in the Mid region significantly enhanced the corresponding Mid:ON ratios of these three strong LTRs, alluding to augmented gene expression noise. For instance, the Mid:ON ratios of 4-κB LTR were 0.07 ± 0.01 and 0.23 ± 0.01 in the absence and presence of MG-132, respectively (Fig 4E). Thus, our data confirm that the LTRs containing more copies of NF-κB motifs demonstrate low-level gene expression noise, which increases significantly when NF-κB signaling is compromised. Percentage cell viability was comparable among all the κB-variant viral strains (Fig 4F) in both treated and non-treated samples. The Tat transcript levels for the stronger LTRs were comparable with or without MG-132 treatment (Fig 4G). We observed a similar profile of augmented gene expression noise following treatment with two other NF-κB pathway inhibitors: Bay 11-7082 (IκBα-degradation inhibitor) and Flavipiridol (IKK inhibitor).

### The gene expression noise profile of LTR-variant viral strains is consistent at a constant intracellular concentration of Tat

In the experiments described above, not only the transcriptional strength of the LTR but also the transcriptional activity of Tat varied since the viral promoter regulated Tat expression. Importantly, it is not clear from these experiments whether the bimodal gene expression profile of HIV-1 LTR could be ascribed to promoter strength differences when the intracellular Tat concentrations also varied concomitantly (Fig 2F, left panel). Additionally, previous studies demonstrated that a positive feedback loop could influence gene expression noise (47–49) (PMID: 12808135, 17510665, 20185727). Therefore, we wanted to examine how varying only the transcriptional strength of the LTR, while keeping the Tat transactivation a constant factor, could influence the noise profile of the LTR-variant viral strains. To this end, we refactored the HIV-1 LTR-Tat circuit to uncouple Tat from the positive feedback loop and supplied Tat from an inducible Tet-ON promoter (Fig 5A). This system modulates intracellular Tat levels concomitant with Dox concentrations. More importantly, the system also permits the evaluation of gene expression of the variant LTRs discordant for the number of NF-κB motifs at a comparable intracellular concentration of Tat. The Tet-ON Tat vector was integrated into Jurkat T-cells to generate a cell line expressing Tat stably. Tat expression was Dox-dependent and directly proportional to Dox concentration without cytotoxicity (Fig 5B). Based on the dose-response, we selected to use Dox at 800 ng/ml to induce Tat expression at the highest level.

**Fig 5:**
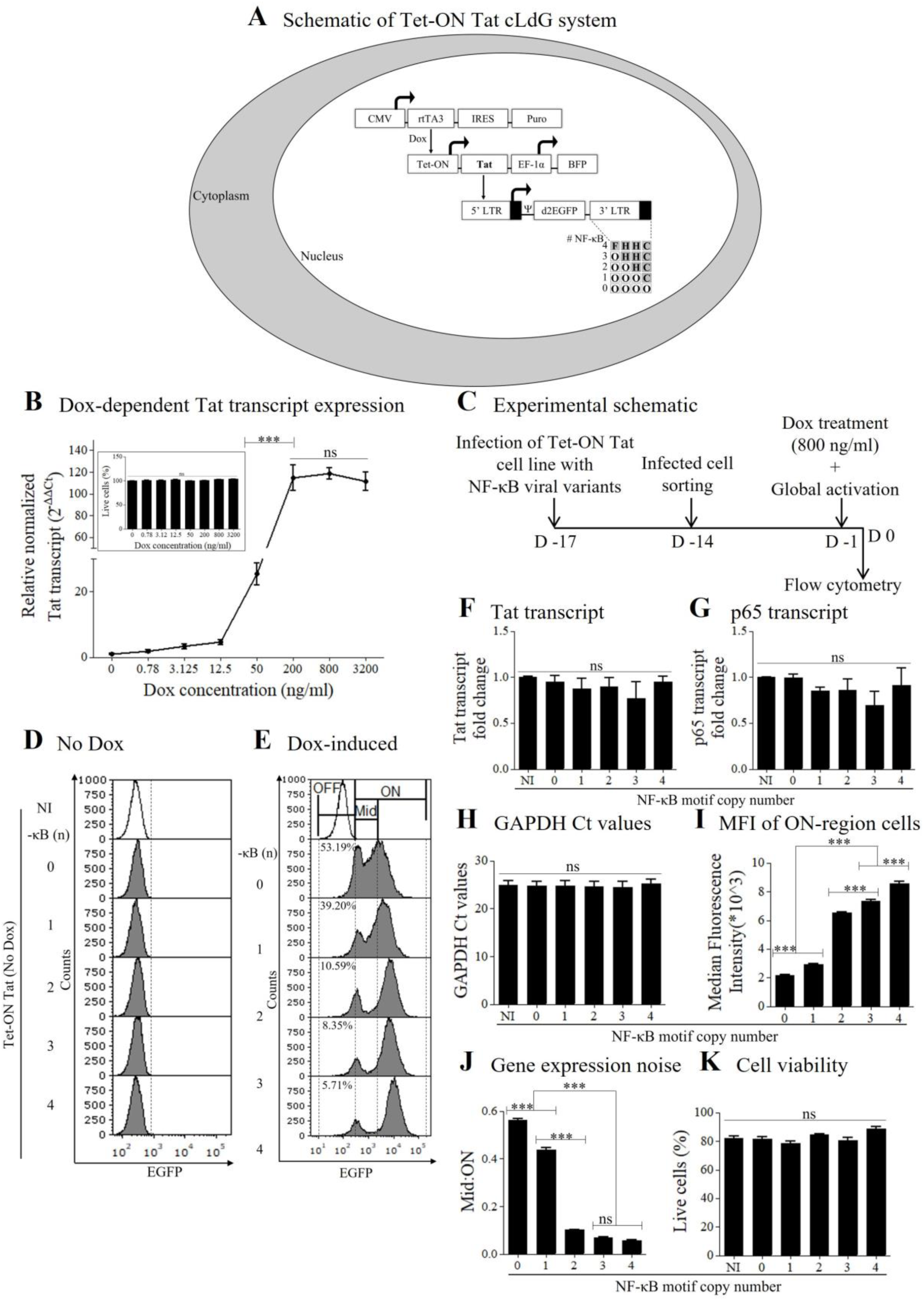
An inverse correlation between the number of NF-κB motifs and gene expression noise of HIV-1C LTR at constant levels of Tat. (A) Schematic representation of the Dox-dependent Tat expression. The stable Tet-ON Tat cell line was infected with the sub-genomic HIV-1C LTR-variant viral strains expressing d2EGFP. (B) Dox-dependent Tat transcript expression. Data represent relative and normalized Tat transcript levels at increasing concentrations of Dox. Mean Tat transcript values from experimental triplicates ± SD are plotted. Inset: Mean percentage values of live cells from experimental triplicates ± SD are plotted. (C) Experimental schematic. Tet-ON Tat cell line was infected with the LdG LTR-variant viral strains, and the infected cells expressing EGFP were sorted. Two weeks following EGFP relaxation, the cells were treated with Dox (800 ng/ml) and a T-cell activator cocktail for 24 h and analyzed using a flow cytometer. Representative stacked histogram profiles depicting gene expression of the Tet-ON Tat LTR-variant viral strains following activation with a cocktail of T-cell activators in (D) the absence or (E) the presence of Dox (800 ng/ml). The open histogram represents non-infected, activated cells and solid histograms represent infected and activated cells. The percentages of the Mid-region cells for each LTR-variant viral strain are depicted in the histogram plots. NI, no infection. Fold change of (F) Tat, (G) p65 transcripts relative to no infection control, and (H) Absolute GAPDH Ct values across different variant viral strains are depicted. Data represent mean values from three independent experiments ± SEM. One-way ANOVA with Bonferroni’s multiple comparison test was performed for statistical evaluation (ns, not significant). NI, no infection. (I) MFI of EGFP^+ve^ cells in the ON region, (J) gene expression noise represented as a Mid:ON ratio, and (K) percentages of activated live cells relative to their no activation control are plotted. Mean values from experimental triplicates ± SD are plotted. Data are representative of three independent experiments. One-way ANOVA with Bonferroni’s multiple comparison test was used for statistical evaluation (***, p<0.0001; ns, not significant). NI, no infection.

We constructed a new panel of subgenomic reporter viral strains, cLdG, analogous to the cLdGIT LTR panel, containing a varying number of NF-κB motifs in the LTR, ranging from 0 to 4. Only EGFP, but not Tat, was encoded by the viral vectors under the LTR. Jurkat T-cells were infected with the viral strains individually, treated with Doxycycline and a cocktail of T-cell activators. The infected cells were flow-sorted, as depicted schematically (Fig 5C), and were allowed to expand in the absence of activation for two weeks.

In the presence of a cocktail of global activators and without Dox-induction, EGFP expression was not observed from the stable-cell pools infected with the variant viral strains (Fig 5D). When the cells were induced with 800 ng/ml of Dox and activated with the cocktail, a bimodal EGFP expression profile was evident, consistent with the previous reports (50) (PMID: 29045398) (Fig 5E). Importantly, the intracellular concentrations of Tat (Fig 5F) and NF-κB p65 (Fig 5G) transcripts were constant and comparable among the cell pools infected with individual viral strains. GAPDH transcript levels were used as an internal cellular control to validate Tat and p65 transcript expression (Fig 5H). Under these experimental conditions, at a constant intracellular Tat concentration, we found that the mean fluorescence intensities of the cell populations were proportional to the number of NF-κB binding sites in the LTR (Fig 5I). Notably, 24 h after cell activation, the percentages of cells in the Mid region were inversely correlated with the number of NF-κB motifs in the LTR. For instance, 53.18 ± 1.06%, 39.19 ± 1.87%, 10.59 ± 0.23%, 8.35 ± 0.72%, and 5.71 ± 0.05% cells corresponding to 0-, 1-, 2-, 3-, and 4-κB LTRs were present in the Mid region. The Mid:ON ratios of the variant LTRs were also inversely correlated with the NF-κB copy number. The Mid:ON ratios of the three strong LTRs were several folds lower than those of the two weak LTRs, with the difference being significant. For instance, the Mid:ON ratios of 2-, 3-, and 4-κB LTRs were, 0.10 ± 0.002, 0.07 ± 0.003, and 0.06 ± 0.001, respectively, while these values for 0- and 1-κB LTRs were 0.56 ± 0.01 and 0.44 ± 0.01, respectively (Fig 5J). Percentage cell viability was comparable among all the κB-variant viral strains (Fig 5K). Thus, even when the intracellular Tat concentration was normalized, the gene expression noise of the strong LTRs was significantly lower than that of the weak LTRs, consistent with that of variable intracellular levels of Tat (Fig 2G, right panel). These data suggest that under these experimental conditions, the TFBS configuration of the LTR, rather than the transactivation property of Tat, is more crucial in modulating gene expression noise for the viral promoter.

### An increase in the NF-κB motif copy number stabilizes the latent state

Using HIV-1B LTR that contains two genetically identical copies of the NF-κB binding sites in the enhancer region (I and II), Burnett et al. examined how each motif may influence the spontaneous viral rebound out of latency (32) (PMID: 19132086). By mutating one κB-motif at a time, the authors demonstrated that κB-site I, but not κB-site II, is crucial for the spontaneous reactivation of the latent virus. We used an analogous experimental strategy to ask how the number of κB-sites in HIV-1C LTR might influence spontaneous latency reversal. A stronger transcriptional activity of the LTR may be expected to make the latent state relatively unstable, for instance. Alternatively, since gene expression noise of stronger LTRs is significantly low (Figs 2G, right panel, and 3D), the stronger LTRs may establish and maintain latency in a relative stabler form. To understand how an increasing number of NF-κB motifs might impact the stability of the latent state, we measured the spontaneous EGFP expression of latent cell populations belonging to each LTR variant strain of the panel.

Jurkat T-cells harbouring latent proviruses of each LTR-variant viral strain were sorted and expanded as depicted schematically (Fig 6A). Cell populations were allowed to relax and the spontaneous EGFP expression manifested by the cells in the absence of cellular activation was monitored for 72 h at an interval of every 24 h. A panel of representative scatter plots depicting EGFP^+ve^ cells of 1- and 4-κB variant viral strains is presented (Fig 6B), and the percentages of EGFP^+ve^ cells corresponding to all the variant viral strains are plotted across days (Fig 6C). Significantly fewer cells infected with the three viral strains containing stronger LTRs (2-, 3-, and 4-κB) expressed EGFP spontaneously without cellular activation than those of the two weak LTRs (0- and 1-κB). For instance, at 24 h following cell sorting of the latent populations, the percentages of cells rebounding from latency representing the weak LTRs (0-, 1-κB) were 7.37 ± 0.36 and 6.89 ± 0.39, respectively. In contrast, the percentages for the strong LTRs (2-, 3-, and 4-κB) were 2.08 ± 0.11, 1.28 ± 0.06, and 1.36 ± 0.06, respectively, the difference between the strong and weak LTRs being statistically significant (Fig 6C). We also quantitated the intra-cellular Tat transcript levels at 0 h and 72 h time-points (Fig 6D). At 0 h, Tat transcript levels were comparable among the variant viral strains. At 72 h, the Tat-transcript levels increased considerably for all the LTRs; however, the weaker the promoters, the higher was Tat transcript level, broadly agreeing with the percentages of the EGFP^+ve^ cells spontaneously activated (Fig 6C). For instance, Tat-transcript levels were 3- and 2.5-folds higher for 0-κB and 1-κB LTRs, respectively, compared to the 4-κB LTR at 72 h time-point. GAPDH transcript levels were used as an internal cellular control to validate the expression levels of the Tat transcripts (Fig 6E). Thus, these data strongly suggest that the loss of NF-κB sites in the LTR destabilizes the latent state of the virus and increases the basal level activity of the promoter regardless of the activation cues. Our data collectively indicate that the canonical HIV-1C LTRs containing three or four copies of the NF-κB binding motif could establish stable latency, consistent with the low-level gene expression noise of these viral promoters.

**Fig 6:**
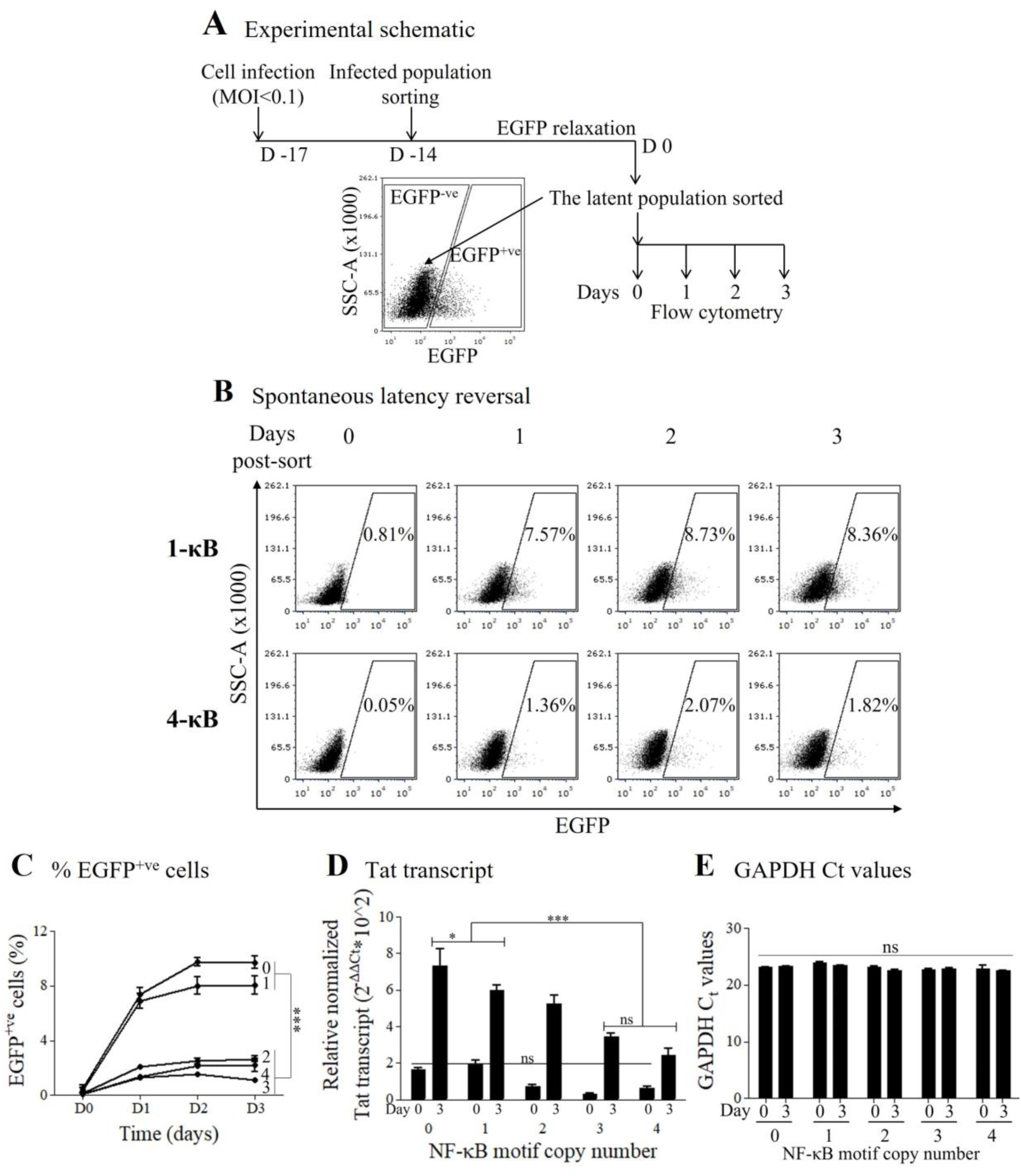
Effect of gene expression noise on HIV-1C latency maintenance. (A) Experimental schematic. Jurkat T-cells were infected with the panel of variant viral strains at an MOI of <0.1. Seventy-two hours following infection, the infected cells were sorted. Following two weeks of relaxation and expansion, EGFP^−ve^ latent cells were sorted, and spontaneous EGFP expression was evaluated every 24 h for 72 h in the absence of cell activation. (B) Representative scatter plots depicting spontaneous EGFP expression of cells infected with 1-κB (top panel) and 4-κB (bottom panel) variant viral strains across days. The percentage of EGFP^+ve^ cells is depicted in each plot. (C) Percentage EGFP^+ve^ cells spontaneously reversing from latency across days are plotted for each strain. Mean percentage of EGFP^+ve^ cells are plotted from three independent experiments ± SEM. Two-way ANOVA with Bonferroni posttest correction was used for statistical evaluation (***, p<0.0001). (D) Relative normalized mean values of Tat mRNA transcripts and (E) Absolute Ct values of GAPDH transcripts at Days 0 and 3 of the κB-variant viral strains from experimental triplicates ± SD are plotted. Data are representative of two independent experiments. Two-way ANOVA followed by Bonferroni’s posttest correction was used for statistical evaluation (***, p<0.0001; *, p<0.05; ns, not significant).

### Strong LTRs demonstrate consistently reduced gene expression noise during latency reversal

Our data demonstrated reduced gene expression noise and maintenance of a stable latent state of strong LTRs compared to weak LTRs. We further examined the profile of gene expression noise during latency reversal. To this end, we activated infected cells with a cocktail of global activators and monitored gene expression at 0, 12, and 24 h (Fig 7A). A panel of representative stacked histograms depicting the temporal EGFP expression pattern of the κB-variant viral strains is presented (Fig 7B), and the kinetics of percentage EGFP^+ve^ cells corresponding to all the variant viral strains are plotted as a function of time (Fig 7C, left panel). Evidently, the efficiency of latency reversal was proportional to the NF-κB motif copy number of the variant viral strains. In other words, the stronger the transcription strength of the LTR, the more efficient the latency reversal. For instance, at 24 h following activation, 40.72 ± 4.96% and 93.56 ± 1.31% cells were activated for 1- and 4-κB variant viral strains, respectively. Additionally, a direct correlation was observed between NF-κB motif copy-number and rapidity of latency reversal among the κB-variant viral strains (Fig 7C, middle panel). The MFI values for 0-, 1-, 2-, 3-, and 4-κB variant viral strains were 3,365.22 ± 1,544.74, 4,482.23 ± 2,047.20, 13,817.50 ± 1,295.51, 16,025.65 ± 2,265.62, and 18,146.05 ± 1,810.30 RFU, respectively, at 24 h of activation.

**Fig 7:**
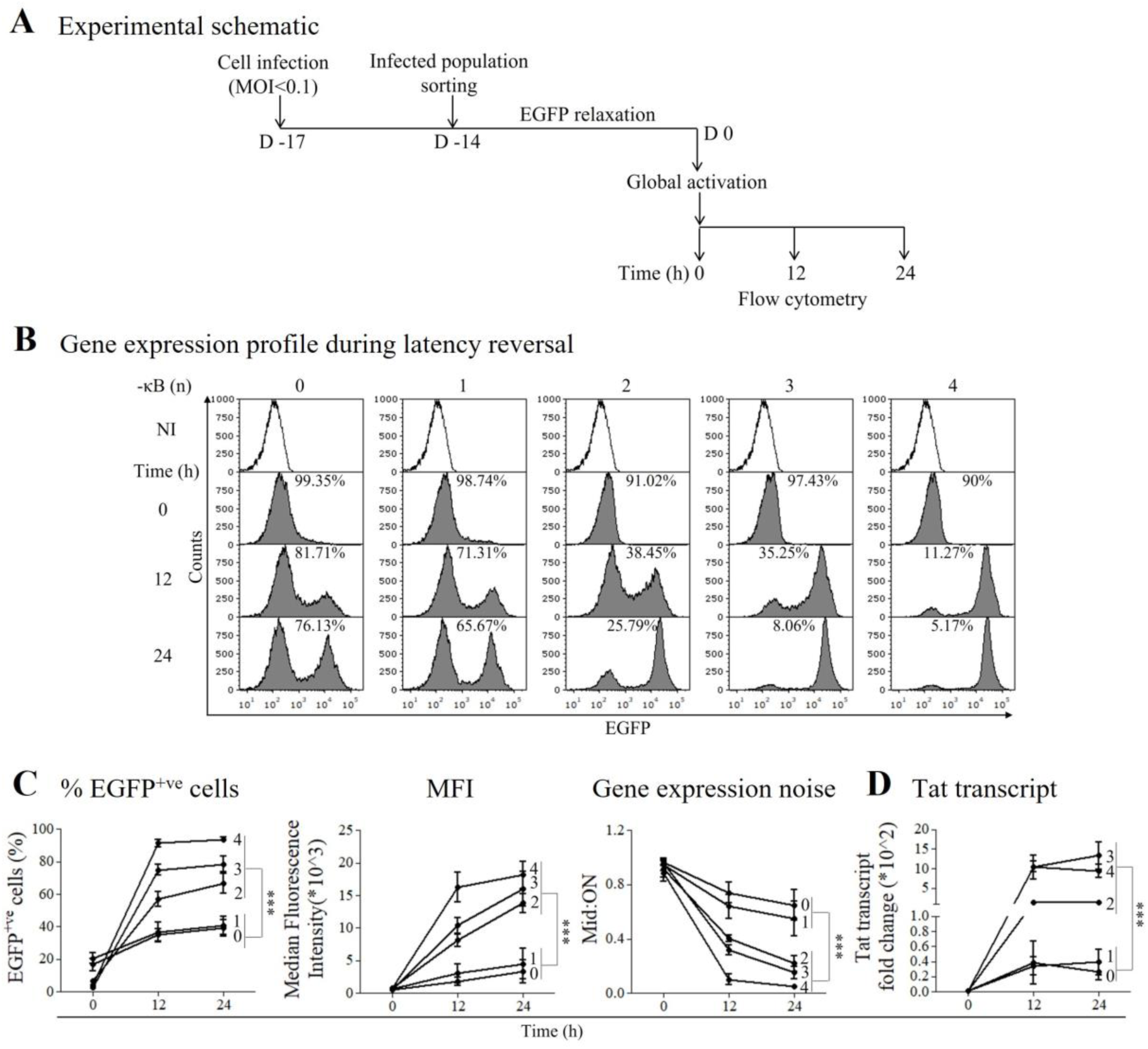
The profile of gene expression noise during latency reversal. (A) Experimental schematic. Jurkat T-cells were infected with the panel of κB-variant viral strains at an MOI of <0.1. Seventy-two hours following infection, the infected cells were sorted expanded for two weeks, activated with a cocktail of global activators, and analyzed at 0, 12, and 24 h following activation. (B) Representative stacked histograms depicting EGFP expression of the variant viral strains across time. The open histogram represents uninfected and activated cells; solid histograms represent infected and activated cells. The percentages of Mid-region cells corresponding to each LTR-variant viral strain across time are depicted. (C) The percentages (left panel), the MFI values (middle panel), and gene expression noise (right panel) of EGFP^+ve^ cells are plotted. Data represent mean values from four independent experiments ± SEM. Two-way ANOVA followed by Bonferroni’s posttest correction was used for statistical evaluation (***, p<0.0001). (D) Fold change values of Tat transcript across time for different variant viral strains are plotted. Data represent mean values from three independent experiments ± SEM. Two-way ANOVA followed by Bonferroni’s posttest correction was used for statistical evaluation (***, p<0.0001).

The percentage of Mid-region cells was consistently low for strong LTRs compared to weak LTRs. Following activation, we measured gene expression noise on a temporal scale by estimating the Mid:ON ratios (Fig 7C, right panel). Gene expression noise was significantly lower for strong LTRs during latency reversal compared to weak LTRs. For instance, the Mid:ON ratio of the 4-κB variant strain at 12 h and 24 h were 0.10 ± 0.04 and 0.05 ± 0.004, respectively, whereas for the 1-κB variant, these values were 0.64 ± 0.08 and 0.55 ± 0.11, respectively. The intracellular Tat concentrations increased as a function of time and NF-κB copy-number, in agreement with the percentages of EGFP^+ve^ cells (Fig 7D). Thus, we conclude that strong LTRs demonstrate efficient and rapid latency reversal compared to weak LTRs. Additionally, gene expression noise was consistently low for strong LTRs during latency reversal.

### Contribution of NF-κB motif genetic variation to gene expression noise of the LTR

HIV-1C LTR contains more copies of the NF-κB motif in the enhancer region than other subtypes (34) (PMID: 23132857). Importantly, these additional κB-motifs are also genetically distinct from the two copies of the genetically identical κB-motif, called the H-κB site (GGGACTTTCC). The C-κB site (GGG<u>G</u>C<u>G</u>TTCC) proximal to the Sp-1 III motif and the F-κB site (GGGACTTTC<u>T</u>) proximal to the upstream RBEIII motif are genetically distinct from the H-κB site. Thus, the NF-κB motif profile of HIV-1C is characterized by a variation not only in the number of these motifs but also in the genetic variation of the additional motifs. We constructed a panel of HIV-1C LTR to examine how the genetically distinct κB-motifs individually might contribute to gene expression noise of the viral promoter when these motifs are arranged tandemly in a cluster. Using the canonical HHC-LTR containing three copies of the NF-κB motif (two and one copies of the H-κB and C-κB motifs, respectively, in that order), a panel of three LTR-variant viral strains was generated containing homotypic clusters of each type of the NF-κB motifs: HHH, FFF, and CCC (Fig 8A). We asked whether genetically distinct κB-motifs could modulate gene expression noise differentially when present in multiple copies. We performed gene expression analysis of the variant viral strains following infection and activation using flow cytometry (Fig 8B).

**Fig 8:**
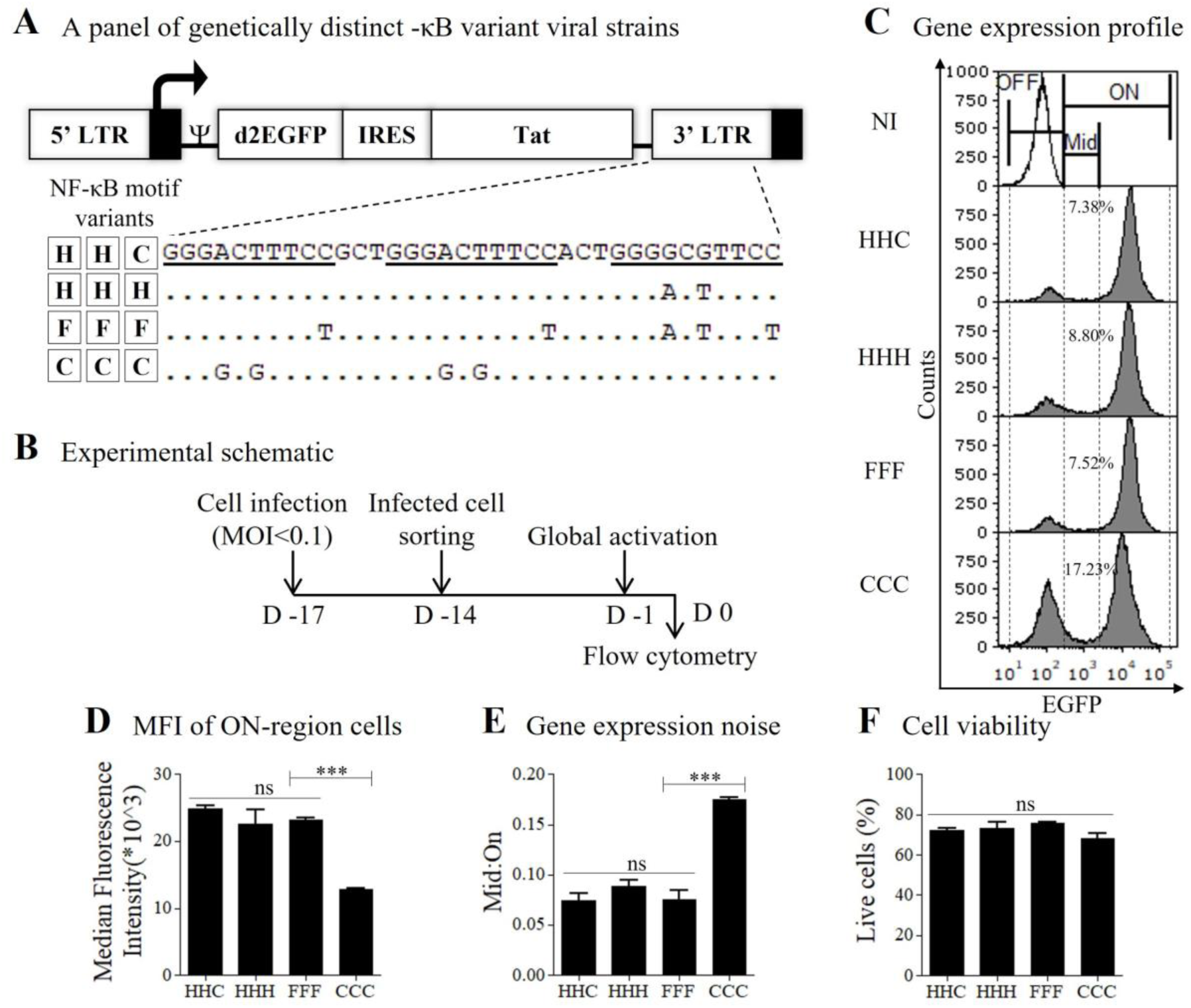
Contribution of genetically distinct NF-κB motifs to gene expression noise. (A) A panel of three cLdGIT sub-genomic variant strains containing genetically distinct homotypic NF-κB binding motifs (HHH, FFF, CCC) derived from the canonical HHC-LTR. The nucleotide sequences of the NF-κB motifs are underlined. Dots represent sequence identity. (B) Experimental schematic. Jurkat T-cells were infected with different variant viral strains at an MOI of <0.1. After 72 h following the infection, the cells were sorted using a flow sorter. Following two weeks of EGFP relaxation and expansion, cells were activated for 24 h with a cocktail of T-cell activators (TNF-α, PMA, HMBA), stained for cell viability, and analyzed using flow cytometry. (C) Representative stacked histogram profiles. The dotted line delineates the OFF, Mid, and ON regions. The open histogram represents non-infected and activated cells; solid histograms represent infected and activated cells. The percentages of Mid-region cells for each NF-κB genetic variant viral strain are depicted in the histogram plots. NI, no infection. (D) MFI of EGFP^+ve^ cells in the ON region, (E) the Mid:ON ratio as a measure of gene expression noise, and (F) the percentage of activated live cells relative to their no activation control for each variant viral strain are plotted. Data represent mean values from three independent experiments ± SEM. One-way ANOVA followed by Bonferroni’s multiple comparison test was used for statistical evaluation (***, p<0.0001; ns, not significant).

A histogram depicting the gene expression profile of the homotypic LTR variants is presented (Fig 8C). The MFI of the canonical HHC-LTR was 24,869.18 ± 209.14 RFU (Fig 8D). The homotypic LTRs FFF and HHH demonstrated MFI values comparable to those of the canonical promoter, 23,221.52 ± 96.49 and 22,611.43 ± 1,006.90 RFU, respectively. Notably, the MFI of CCC-LTR was approximately half (12,836.55 ± 65.12 RFU) of the HHC-LTR, the difference being statistically significant, suggesting a compromised transcriptional strength of the CCC-LTR. The corresponding Mid:ON ratio values of HHC, HHH, FFF, and CCC were 0.07 ± 0.007, 0.09 ± 0.006, 0.08 ± 0.009, and 0.18 ± 0.002, respectively (Fig 8E). While the Mid:ON ratios of HHC-, HHH-, and FFF-LTRs were comparable, that of the CCC-LTR variant was nearly 2-folds higher. The percentage of cell viability was comparable among all the variant viral strains (Fig 8F). These data are suggestive that the C-κB motif contains characteristically reduced transcriptional strength and significantly augmented gene expression noise among all the genetic variants of the NF-κB motif observed in HIV-1C. High gene expression noise of the C-κB motif also explains why the 1-κB variant viral strain with only the C-κB motif behaved like the 0-κB variant (Fig 2G, right panel).

### Strong viral LTRs demonstrate cooperativity in gene expression

Homotypic clusters of degenerate TFBS of a specific transcription family represent key elements of promoters and enhancers of the human genome and constitute nearly 2% size of the genome (51–53) (PMID: 16670430, 18256240, 20363979). Homotypic clusters of TFBS increase the local concentration of TFs around the regulatory sites (54) (PMID: 14692768), provide functional redundancy (55,56) (PMID: 12459721, 11875036), and most importantly, may favor cooperative binding of TFs to such sites (57) (PMID: 9159075).

The canonical HIV-1B LTR contains two genetically identical H-κB motifs separated by a spacer sequence of four base pairs. The distance between the two motifs is not optimal for cooperative binding of NF-κB, as was experimentally demonstrated (58) (PMID: 19683540). In contrast, the canonical 3- and 4-κB LTRs of HIV-1C not only contain a larger number of the κB-motifs in the LTR, but the spacer length separating the motifs is only three base pairs, closer to the optimal distance of two base pairs estimated optimal for cooperative binding of NF-κB (see discussion) (34,36) (PMID: 23132857, 34899661). Given these important variations in the organization of the homotypic clusters of NF-κB motifs in HIV-1C LTR, we wanted to understand if some level of cooperative binding would be possible in HIV-1C. Cooperativity in gene expression of a promoter could be determined by estimating the H value from the gene expression trajectory. A value of H=1 represents the independent binding of TFs to the cognate site and the absence of cooperativity, H>1 indicates positive cooperativity where the binding of one TF facilitates the direct/indirect binding of other TFs to the TFBS, and H<1 signifies the lack of cooperativity.

To this end, we infected Jurkat T-cells with strong (2-, 3-, and 4-κB LTRs) or weak (1-κB) LTR-variant viral strains. Following infection, the cells were sorted and expanded for two weeks. The latently infected stable polyclonal population of cells was sorted and activated using a cocktail of T-cell activators. The fluorescence of the cells was monitored for 22 h at two-hour intervals using flow cytometry (Fig 9A). We observed two different trajectories of gene expression kinetics of the strong and the weak LTRs. The strong LTRs demonstrated a sigmoidal EGFP trajectory, whereas the weak LTR demonstrated relatively linear kinetics (Fig 9B). The H values of the 4-, 3-, 2-, and 1-κB LTR were 3.75, 3.56, 2.61, and 1.38, respectively, alluding to the cooperative binding of NF-κB to the homotypic clusters of HIV-1C LTRs containing more than one NF-κB motif. The nature of the cooperativity, whether direct or indirect, warrants further investigation.

**Fig 9:**
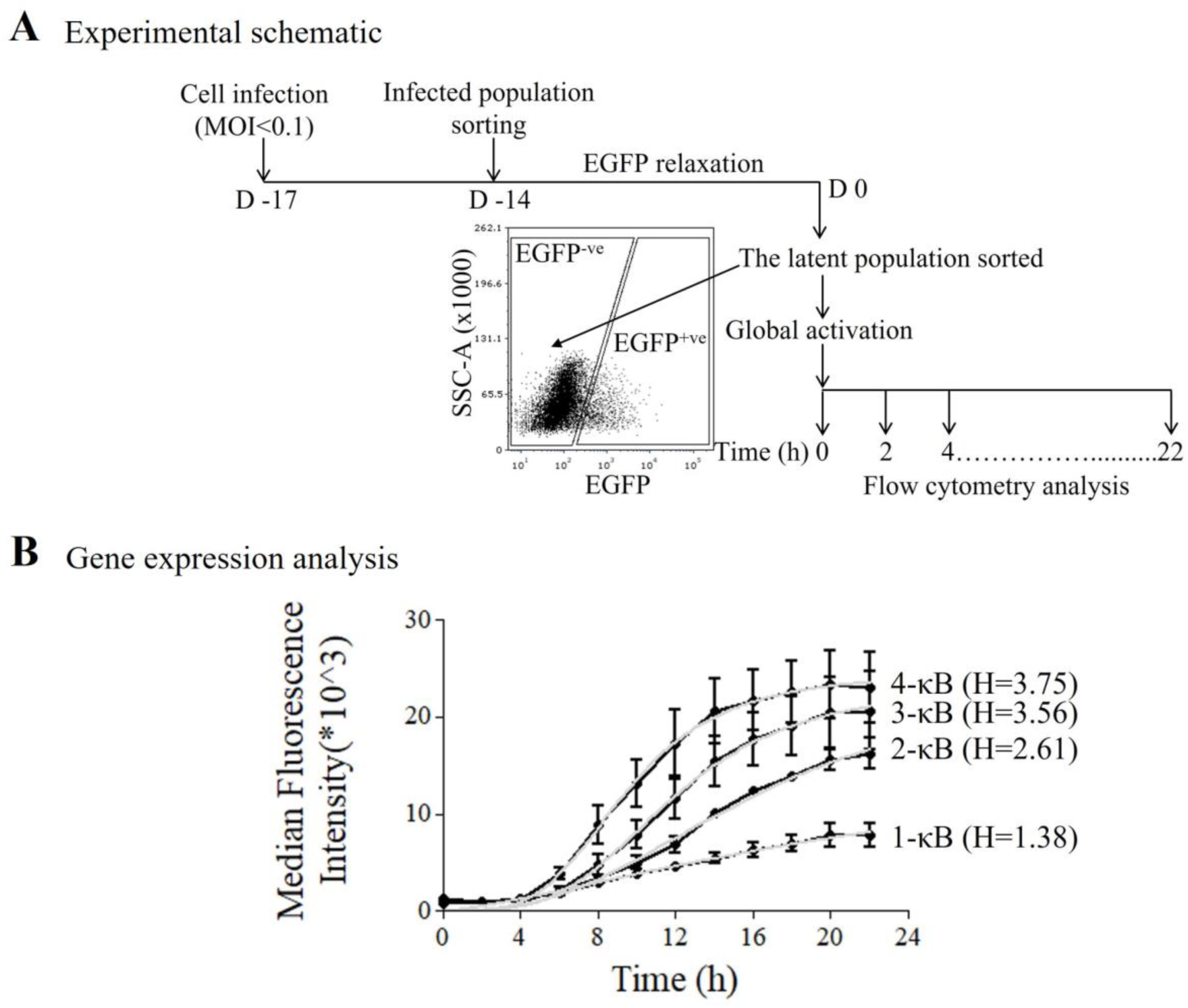
Positive cooperativity in strong HIV-1C LTR. (A) Experimental schematic. Jurkat T-cells were infected with the strong (2-, 3-, and 4-κB) and weak (1-κB) LTR-variant viral strains. The cells were sorted 72 h following the infection, allowed to relax and expand for ~two weeks, and the latent population was sorted. The latent cells were activated with a cocktail of T-cell activators and analyzed for EGFP expression at an interval of 2 h for 22 h using flow cytometry. (B) Kinetic profiles of gene expression of 1-, 2-, 3- and 4-κB variant viral strains following activation. Data represent mean MFI values ± SEM from three independent experiments. Hill coefficient (H) was calculated by fitting the MFI values in a non-linear Hill equation curve. The grey line represents the best fit non-linear curve, and the black line represents the experimental data.

## Discussion

In the present work, we attempted to examine the effects of the enhanced transcriptional strength of HIV-1C LTR on gene expression noise. Stochastic fluctuations in Tat levels are believed to influence the choice between active replication and the transcriptionally silent state of HIV-1 (14,20) (PMID: 16051143, 18344999). Since the LTR directly controls Tat expression, the enhanced transcriptional strength of the former is expected to increase the intracellular concentration of Tat. Further, given that Tat positive feedback amplifies the fluctuations in gene expression to drive phenotypic variability (14) (PMID: 16051143), it was necessary to evaluate how enhanced gene expression of the viral promoter influences transcriptional noise; therefore, the ON/OFF decision-making of the virus.

Among stochastic models, no consensus exists regarding how gene expression noise correlates with transcriptional strength and the number of TFBS in various biological groups. A few studies in yeast (25,26) (PMID: 15166317, 23565060), and mammalian cells (24) (PMID: 20159560) demonstrated that an enhanced transcriptional activity or increased TFBS copy number leads to a concomitant reduction in the magnitude of gene expression noise; while a few other studies reported results contrary to this notion (27–29,31,48) (PMID: 16715097, 16699522, 17048983, 20185727, 25030889). All these studies collectively demonstrated that gene expression noise is a function of the features of a promoter architecture such as the number and strength of TFBS, nucleosome occupancy and positioning, strength of TATA box, etc.

### NF-κB motif cooperativity

HIV-1 LTR presents a complex promoter architecture where several TFBS, especially the NF-κB motifs, are arranged tandemly and in proximity. For example, in HIV-1B LTR, the enhancer comprises two genetically identical copies of the H-κB motif, separated by a spacer of four base pairs. Diverse experimental strategies demonstrated the absence of cooperativity between the two NF-κB binding motifs (58,59) (PMID: 11970949, 19683540). Non-cooperative functioning is also typical of several inducible cellular promoters containing multiple copies of the NF-κB motif, even up to six (24) (PMID: 20159560). The NF-κB motifs in such clusters of cellular promoters are not positioned in proximity, unlike in HIV-1 LTR. The authors demonstrated that the non-cooperative functioning of clustered NF-κB motifs is crucial in providing transcriptional output proportional to the strength of inducing a signal in a graded manner. Importantly, the spacer sequence length separating two such motifs appear critical in determining the nature of the interaction between NF-κB motifs in a cluster. When the distance between the adjacent NF-κB motifs of the *NFKBIA* cellular promoter was reduced, using simulation, the graded response transformed into a binary response suggestive of cooperativity between the κB-motifs. The authors propose that cellular promoters, by maintaining the mean distance between two NF-κB binding sites not less than 23 bases, ensure a graded response proportional to the signal input, in the absence of cooperativity. Additionally, a larger number of NF-κB binding sites are believed to make cellular promoters more sensitive to the nuclear concentration of NF-κB.

In the light of the above proposition, it is noteworthy that the NF-κB motif cluster of HIV-1C could be distinct from that of other HIV-1 subtypes given three specific characteristics. First, HIV-1C LTR typically contains a larger number of NF-κB binding sites, three (HHC) or four (FHHC), than that of other subtypes that contain only one or two (HH). Second, while the two NF-κB motifs of HIV-1B LTR are genetically identical, the HHC-LTR of HIV-1C contains two genetically distinct motifs (H and C), and the FHHC-LTR contains three distinct motifs (H, C, and F). Sequence variations of NF-κB binding motifs could induce differential conformational changes in the p50-p65 heterodimer recruited, probably with comparable affinity, which in turn could cause the assembly of enhanceosomes of different host factor composition, thus leading to differential outcomes (59) (PMID: 11970949). Lastly, the spacer sequence separating the NF-κB binding sites in the HHC-LTR contains only three base pairs compared to four in the HH-LTR of HIV-1B. Of note, the length of an optimal spacer separating two NF-κB binding sites was estimated to be two base pairs to permit cooperativity in the LTR (41,58) (PMID: 19683540, 17869269). With a spacer sequence comprising one base pair short, the HIV-1C NF-κB cluster is better positioned for possible cooperativity. Thus, with a larger number of NF-κB binding motifs present in the cluster, the genetically diverse NF-κB motifs possibly assembling enhanceosomes of diverse composition, and a shorter spacer sequence separating the proximal motifs, all may permit cooperativity among NF-κB binding motifs. Additionally, the length of NF-κB binding sequences could be variable, occasionally comprising 11 base pairs (60–62) (PMID: 9450761, 10360983, 12048232). If any of the NF-κB motifs in HIV-1C LTR comprises 11 base pairs, the spacer sequence length will be reduced to only two base pairs, creating an ideal context that could permit cooperativity. Notably, the sigmoidal curve of gene expression characterized by H>1 manifested by the strong LTRs (2-, 3- and 4-κB LTRs) compared to a linear curve of H~1 of the weak promoter (1-κB LTR) alludes to the presence of cooperativity among the adjacent NF-κB motifs (Fig 9B). Additionally, efficient and rapid viral latency reversal of strong promoters in comparison with weak promoters alludes to the presence of cooperativity among the multiple κB-motifs (Fig 7C, left and middle panel).

### The C-κB motif appears to be transcriptionally noisier

The C-κB motif is unique for HIV-1C LTR and is positioned at the crucial spot adjacent to the Sp-1III site displacing the canonical H-κB motif from this location. We previously demonstrated the co-evolution of both the centrally located C-κB motif and the HIV-1C-specific Sp-1III site (40) (PMID: 27194770). Importantly, the HIV-1C-specific Sp-1III site can function in association with the C-κB motif but not with the H-κB motif. That a genetically distinct C-κB motif is positioned in the crucial position of the viral promoter, we wanted to ask how this novel κB-motif might influence the transcriptional noise of the viral promoter. The engineered LTR containing a homotypic cluster of three C-κB motifs (CCC-LTR) demonstrated significantly higher gene expression noise than the FFF-, HHH- or the canonical HHC-LTR (Fig 8E). Additionally, we also observe high gene expression noise in 0- (mutations in all the κB-motifs) and 1-κB (bearing the C-κB motif) promoters (Figs 2G, right panel and 3D).

These data allude to the essential nature of the C-κB motif to the functioning of HIV-1C LTR, even though the latter causes high levels of gene expression noise, which is precluded by the presence of the two canonical H-κB motifs. Thus, the C-κB motif appears noisier than the H- and F-κB motifs. The indispensable nature of the C-κB motif to the HIV-1C is evident that the vast majority of the viral sequences belonging to this viral subtype deposited in the extant databases invariably contain this motif. Furthermore, the gradual disappearance of the 4-κB (FHHC) LTR configuration and its possible replacement by a 3-κB (FHC) LTR are suggestive of HIV-1C attempting to optimize transcriptional noise levels. The invariable presence of the C-κB motif in HIV-1C, despite the significantly higher-level transcriptional noise it causes, is indicative of a crucial function the C-κB motif, but not the H-κB motif, provides to HIV-1C.

### The significance of reduced gene expression noise to HIV-1 latency maintenance

We altered the transcriptional strength of the LTR by varying the number of NF-κB motifs and attempted to understand the effect of modulated gene expression noise on maintaining a stable latent state in the absence of cellular activation (Fig 6). Strikingly, the LTRs containing at least two functional NF-κB motifs (2-, 3-, and 4-κB LTRs) showed significantly low levels of spontaneous latency reversal, unlike the two weak promoters (0- and 1-κB LTRs). The maintenance of stable latency by stronger LTRs may be ascribed to kinetically steady recruitment and maintenance of transcription-suppressing p50:p50 homodimers (63) (PMID: 16319923). Tandemly arranged, degenerate TFBS are evolutionarily conserved in the human genome, and such elements increase the binding stability of TFs (51) (PMID: 16670430). Additionally, the same NF-κB binding sites are shared by NFAT family proteins in a mutually exclusive manner (64,65) (PMID: 12949493, 18462673). Further, we previously demonstrated that NFAT1 predominantly binds the latent promoter (39) (PMID: 32669338). Thus, the spontaneous latency reversal and high basal-level activity of the weak LTRs compared to the strong LTRs could be ascribed to stable recruitment of repressive transcription complexes to the viral promoter, as proposed schematically (Fig 10). We propose that multiple copies of the NF-κB motif in the enhancer region of the LTR ensure efficient recruitment of transcription promoting or suppressing complexes stably, in the presence or absence of activation, respectively.

**Fig 10:**
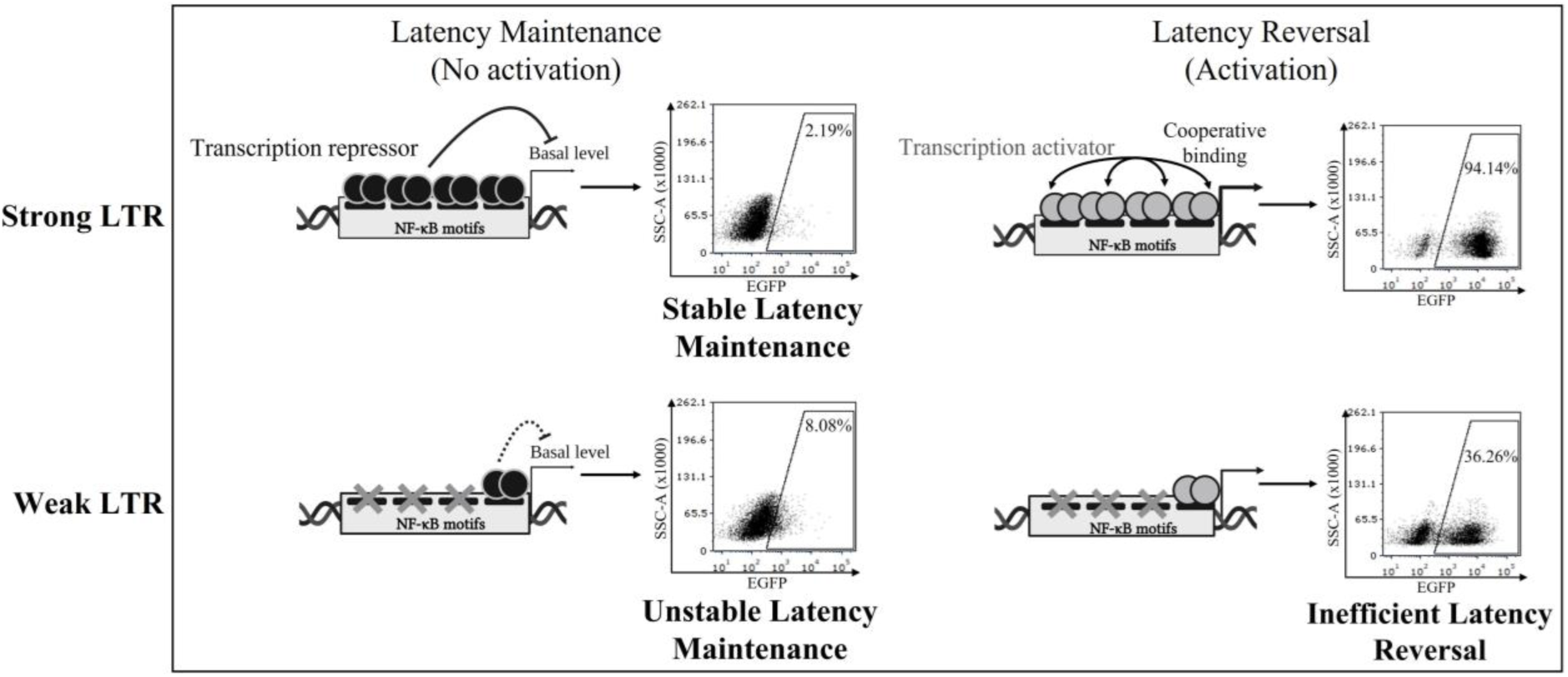
A schematic model depicting the probable influence of multiple NF-κB motifs on latency maintenance and reversal. The multiple NF-κB motifs of a strong LTR can recruit and maintain a higher concentration of transcription repressors around the LTR, thus exerting a stronger transcription suppression and ensuring low-level transcriptional noise in the absence of optimal activation conditions, ensuring stronger gene expression (top-left panel). Copy number and sequence variations of the NF-κB motif (see Fig 1A) and those of other TFBS in HIV-1C LTR (36) (PMID: 34899661) may contribute to cooperativity. Under the conditions of optimal activation, the multiple κB-motifs of a strong LTR recruit transcription activator complexes through direct and/or indirect cooperativity, efficiently reverse viral latency, and maintain stronger levels of viral transcription (top-right panel). In contrast, a weak LTR lacking multiple NF-κB motifs cannot maintain avid viral latency due to sparse recruitment of transcription repressor complexes, thus leading to high levels of transcriptional noise (bottom-left panel). Under the conditions of activation, a weak LTR can maintain only low-level gene expression (bottom-right panel). The data presented in the present work are consistent with this model.

### Implications for evolutionary selection in HIV-1C

The outcome that enhanced transcriptional strength is inversely correlated with transcriptional noise, although anticipated, is surprising since gene expression noise of the LTR is expected to play a crucial role in the ON/OFF decision-making (14,20) (PMID: 16051143, 18344999). The cost of reduced transcriptional noise of the LTR, the consequence of the enhanced transcriptional strength, may lead to the loss of replication fitness of viral strains that contain more NF-κB motifs in the enhancer, as exemplified by our recent report (36) (PMID: 34899661). The proportion of 4-κB viral strains was found to be 1-2% (n= 607 samples) in 2003 in India (38) (PMID: 15184461), which increased to 20-30% (n= 159 samples) in ten years (34). However, in a recent clinical study conducted at four clinical sites spanning the country, we found that the proportion of the 4-κB viral strains dropped to approximately 6% (n= 455 samples), after approximately twenty years since the first report (36) (PMID: 34899661). Thus, the LTR-variant viral strains endowed with transcriptional activity higher than necessary appear unable to sustain long-term replication fitness.

In this context, the emergence of a new LTR of HIV-1C containing only three NF-κB motifs like the canonical HIV-1C LTR is noteworthy. However, unlike in the canonical HIV-1C LTR containing only two types of κB-motifs (HHC-LTR), in the emerging FHC-LTR, all the three κB-motifs are genetically variant (36) (PMID: 34899661). Given that the F-κB motif is known to be associated only with the 4-κB (FHHC) LTR and that the proportion of the 4-κB containing viral strains is decreasing, it appears plausible that the FHHC viral strains relinquish one of the two H-κB motifs to regain an LTR configuration containing only three NF-κB binding motifs; however, without sacrificing the genetic diversity of the NF-κB binding motifs.

Furthermore, several other promoter-variant strains of HIV-1C co-duplicate an NF-κB motif and an RBEIII site with or without the TCF-α/LEF sequence (36) (PMID: 34899661). Especially the RBEIII motif can mediate the coordinated binding of RBF-2 (66) (PMID: 21813151) and AP-1 factors. The constitutive components of RBF-2 and AP-1 can suppress transcription, especially in HIV-1 (67,68) (PMID: 22037610, 23236059). Thus, several promoter-variant strains have been emerging in HIV-1C, formulating diverse TFBS profiles.

The evolution of HIV-1C LTR appears to be directional, moving towards enhanced transcriptional strength under the conditions of cell activation and establishing and maintaining avid latency when not activated, all without augmenting transcriptional noise beyond a certain threshold level.

Collectively, our data allude to the need to maintain a delicate balance between transcriptional strength and gene expression noise in the functioning of the LTR. Since the ‘OFF’ state is the default monostable state of the virus (69,70) (PMID: 12682019, 17194214), modulation of gene expression noise is expected to have severe consequences for viral latency decisions.

Our data raise an important question that was beyond the scope of the present study. How the observed TFBS profile variation is likely to influence the replication fitness of several emerging LTR-variant viral strains that we reported recently needs evaluation. We acknowledge a few technical limitations of the present study. Our attempts to evaluate gene expression noise using full-length HIV-1C strains were unsuccessful owing to high cytopathic effects in the infected cells. We employed a destabilized form of EGFP (d2EGFP) with a half-life of ~two h for examining the stochastic processes of gene expression. However, such stochastic processes may occur on scales of seconds, which reporter proteins fail to capture.

## Acknowledgments

Several reagents were obtained through the NIH HIV Reagent Program, Division of AIDS, NIAID, NIH. We thank Prof. Hemalatha Balaram (JNCASR, India) for the intellectual discussions. We thank Dr. Malini Menon and Chhavi Saini (JNCASR, India) for the Tet-ON Tat cell line.

## Ethics Statement

The studies were approved by the Institutional Biosafety Committee of JNCASR. The Human Ethics and Biosafety Committee of JNCASR approved the collection of blood from healthy controls.

## Author contributions

SP, conception and design, acquisition of data, analysis, and interpretation of data, drafting or revising the article; VJ, acquisition, and analysis of data; NN, acquisition of data; UR, conception and design, funding acquisition, validation, reviewing, and editing the article.

## Competing Interests

We declare that no competing interests exist.

## Notes

### Competing Interest Statement

The authors have declared no competing interest.

